# Plant pathogenic bacterium can rapidly evolve tolerance to an antimicrobial plant allelochemical

**DOI:** 10.1101/2021.05.21.445234

**Authors:** Carrie Louise Alderley, Samuel Terrence Edwards Greenrod, Ville-Petri Friman

## Abstract

Crop losses to plant pathogens are a growing threat to global food security and more effective control strategies are urgently required. Biofumigation, an agricultural technique where *Brassica* plant tissues are mulched into soils to release antimicrobial plant allelochemicals called isothiocyanates (ITCs), has been proposed as an environmentally friendly alternative to agrochemicals. While biofumigation has been shown to suppress a range of plant pathogens, its effects on plant pathogenic bacteria remain largely unexplored. Here we used a laboratory model system to compare the efficacy of different types of ITCs against *Ralstonia solanacearum* plant bacterial pathogen. Additionally, we evaluated the potential for ITC-tolerance evolution under high, intermediate and low transfer frequency ITC exposure treatments. We found that allyl-ITC was the most efficient compound at suppressing *R. solanacearum* growth, and its efficacy was not improved when combined with other types of ITCs. Despite consistent pathogen growth suppression, ITC tolerance evolution was observed in the low transfer frequency exposure treatment, leading to cross-tolerance to ampicillin beta-lactam antibiotic. Mechanistically, tolerance was linked to insertion sequence movement at four positions in genes that were potentially associated with stress responses (H-NS histone like protein), cell growth and competitiveness (acyltransferase), iron storage ((2-Fe-2S)-binding protein) and calcium ion sequestration (calcium-binding protein). Interestingly, pathogen adaptation to the growth media also indirectly selected for increased ITC tolerance through potential adaptations linked with metabolism and antibiotic resistance (dehydrogenase-like protein) and transmembrane protein movement (Tat pathway signal protein). Together, our results suggest that *R. solanacearum* can rapidly evolve tolerance to allyl-ITC plant allelochemical which could constrain the long-term efficiency of biofumigation biocontrol and potentially shape pathogen evolution with plants.

## Introduction

Plant pathogens are a growing threat to global food security, accounting for up to 40% of crop losses annually (Savary et al., 2012). The phasing out of environmentally toxic chemical fumigants, such as methyl bromide, has directed attention towards alternative biocontrol strategies (Qin et al., 2004). Plant-derived antimicrobial allelochemicals, such as phenolic acids, terpenes and volatile isothiocyanates (ITCs), are naturally exuded by the roots of legumes (Mondal et al., 2015; Wink, 2013), cereals (Larkin & Halloran, 2015; Mazzola & Gu, 2002) and other crops such as *Brassica* (Kirkegaard et al., 1996; Sarwar et al., 1998). These compounds could potentially be used to control pathogens by biofumigation, which involves mulching plant tissues into soils to release biocidal allelochemicals. While biofumigation has previously been shown to suppress the growth of soil-borne fungal (Angus et al., 1994; Rumberger & Marschner, 2003; Sarwar et al., 1998), nematode (Lord et al., 2011; Ngala et al., 2015) and bacterial pathogens (Hu et al., 2015; Ji et al., 2007), outcomes are still varied, ranging from clear pathogen suppression (Larkin & Griffin, 2007; Matthiessen & Kirkegaard, 2006) to having no effect (Hartz et al., 2005; Kirkegaard et al., 2000; Stirling & Stirling, 2003). A better understanding of the antimicrobial and biocidal effects of plant allelochemicals on pathogens is thus required.

The success of biofumigation is influenced by various factors including soil conditions, the biofumigant plant species, timing of application and the half-life of biocidal compounds (Matthiessen & Kirkegaard, 2006). The biocidal effects of *Brassica*-based biofumigation are believed to result primarily from the release of toxic ITCs from their glucosinolate (GSL) pre-cursors (Gimsing & Kirkegaard, 2009; Lord *et al*., 2011; Matthiessen & Kirkegaard, 2006). Moreover, other allelochemicals such as dimethyl sulfide and methyl iodide might contribute to the biocidal activity of biofumigant plants (Vervoort et al., 2014; Wang et al., 2009). Even though ITC-liberating GSL levels can potentially reach as high as 45.3 mM/m^2^ following initial mulching of plant material into the soil (Kirkegaard & Sarwar, 1998), their concentrations often decline rapidly due to high volatility, sorption to organic matter, leaching from the soil and microbial degradation (Frick et al., 1998; Gimsing et al., 2007; Hanschen et al., 2015; Matthiessen & Kirkegaard, 2006; Warton et al., 2001). As ITCs often have short half-lives of up to sixty hours (Borek et al., 1995; Gimsing & Kirkegaard, 2006), it is important to identify ITCs that are highly effective against pathogens even during short-term exposure.

The antimicrobial activity of different types of ITCs can vary depending on their mode of action and the species and genotype of the target pathogen. In the case of bacterial pathogens, several antimicrobial mechanisms have been suggested. For instance, ITCs could damage the outer cell membrane of Gram-negative bacteria leading to changes in cell membrane potential (Sofrata et al., 2011) and leakage of cell metabolites (Lin et al., 2000). Further, it has been suggested that ITCs could bind to bacterial enzymes, such as thioredoxin reductases and acetate kinases and disrupt their tertiary structure and functioning (Luciano & Holley, 2009). It is also possible that some ITCs, such as allyl-ITC, could have multiple targets, making them relatively more toxic to pathogenic bacteria (Luciano & Holley, 2009). However, antimicrobial activity and potential tolerance evolution to ITCs are still poorly understood in plant pathogenic bacteria.

Antibiosis is an important mechanism underlying bacterial competition in soils and soil bacteria often produce and are resistant to several antimicrobials, enabling them to outcompete surrounding bacteria for space and nutrients (Hibbing et al., 2010). Antimicrobial tolerance is also important for plant-bacteria interactions, as it can help bacteria to tolerate antimicrobials secreted by plants, such as coumarins, giving them a selective advantage in the plant rhizosphere microbiome (Stringlis et al., 2018). Such tolerance has recently been shown to evolve *de novo* in *Pseudomonas protegens* CHA0 bacterium against the antimicrobial scopoletin secreted by *Arabidopsis thaliana* (Li et al., 2020). Prolonged exposure to plant allelochemicals could thus select for more tolerant plant pathogen genotypes also during biofumigation and will likely be affected by the strength and duration of ITC exposure, which is important in determining whether potential tolerance or resistance mutations have enough time to sweep through pathogen populations. If the mutations enabling ITC tolerance are costly, their selective benefit could be further reduced by competition or growth trade-offs, leading to loss of tolerance mutations in the absence of ITCs. While ITC concentrations are known to reach antimicrobial levels during biofumigation in the field (Sarwar et al., 1998), no direct experimental evidence for ITC tolerance evolution in plant pathogenic bacteria exists.

To study these questions, we developed a model laboratory system where we tested the growth-inhibiting effects of ITCs produced by Indian mustard (*Brassica juncea*) on *Ralstonia solanacearum* plant pathogenic bacterium, which is the causative agent of bacterial wilt and potato brown rot diseases and a globally important pathogen, affecting over 200 different plant species including various important crops (Elphinstone, 2005; Yabuuchi et al., 1995). The *R. solanacearum* genome is bipartite, consisting of a chromosome and megaplasmid (Salanoubat et al., 2002). Disease control techniques such as crop rotation, the use of clean and certified seeds or resistant plant cultivars, have shown only limited success in controlling *R. solanacearum* (Chellemi et al., 1997; Ciampi-Panno et al., 1989; Ramesh et al., 2009). Indian mustard was chosen as a model biofumigant plant due to its well-established allelochemical properties (Bending & Lincoln, 1999; Kirkegaard & Matthiessen, 2005; Mazzola et al., 2015; Sarwar et al., 1998), which are predominantly caused by the release of allyl, sec-butyl and 2-phenylethyl ITCs (Bangarwa et al., 2011; Olivier et al., 1999; Yim et al., 2016). As these ITCs might vary in their biocidal activity, we first tested to what extent they suppress *R. solanacearum* growth when applied either alone or in combination at concentrations relevant to field biofumigation (Gimsing et al., 2007; Hanschen et al., 2012; Kirkegaard & Sarwar, 1998; Matthiessen & Kirkegaard, 2006; Rudolph et al., 2015). Subsequently, we explored whether long-term exposure to the most effective ITC type could select for resistant or more ITC-tolerant pathogens in the lab, and if ITC tolerance is associated with competitive costs or cross-tolerance to other antimicrobials. It was found that allyl-ITC was the most suppressive allelochemical. However, long-term exposure selected for ITC-tolerant pathogen mutants that also had increased cross-tolerance to the beta-lactam antibiotic ampicillin. At the molecular level, adaptations were associated with a few parallel mutations and loss of insertion sequences mainly in the megaplasmid. Together these results suggest that while Indian mustard could be used as a biofumigant plant against *R. solanacearum* due to the high antimicrobial activity of allyl-ITC, its long-term efficacy could be constrained by rapid ITC tolerance evolution.

## Materials and Methods

### (a) Pathogen strain and culture media

We used a *Ralstonia solanacearum* strain (21415687) which was originally isolated from the river Loddon (phylotype II sequevar 1) in the UK as our ancestral pathogen strain (Source: John Elphinstone, Fera Science, 2014). This strain was chosen as river water is the most common environmental source of potato brown rot outbreaks in the UK (Elphinstone *et al*., 1998), and hence highly relevant for UK *R. solanacearum* epidemics. The strain was cultured in CPG broth (1 g casamino acids, 10 g peptone and 5 g glucose per litre of ddH_2_O) for 48 hours at 28 °C to create cryostocks (20% w/v glycerol) that were preserved at -80 °C. CPG was also used as the main growth media in all experiments except for fitness assays, where lysogeny broth (LB: 10 g tryptone, 5 g yeast, 10 g NaCl per litre of ddH_2_O) was also used as a ‘naïve’ growth media to control the effects of *R. solanacearum* adaptation to CPG media during the selection experiment.

### (b) Comparing the effects of different types of ITCs for pathogen suppression

To determine antimicrobial activity of ITCs, we first identified concentrations that caused a significant reduction in *R. solanacearum* growth relative to the no-ITC control treatments. To this end, we conducted short-term growth assays where *R. solanacearum* was exposed to allyl, sec-butyl and 2-phenylethyl ITCs at 63, 125, 250, 500, 1000, 2500 and 5000 μM concentrations in CPG media (Suppl. Fig. 2). For this experiment, *R. solanacearum* was revived from cryostocks by growing with shaking (250 rpm) for 48 hours at 28 °C before normalising bacterial density to an optical density (OD) reading of 0.1 (600 nm; Tecan, Sunrise), equalling ∼10^7^ cells per ml. This method was consistently used to revive and adjust bacterial densities in all growth experiments. *R. solanacearum* was grown in 200 μl CPG media in different ITC concentrations for 148 hours and bacterial densities were measured every 24 hours (OD600 nm). We found that allyl-ITC concentrations as low as 125 μM inhibited *R. solanacearum* growth, while relatively higher concentrations of 500 μM of sec-butyl and 2-phenylethyl ITC were required to inhibit pathogen growth (Suppl. Fig. 2). Based on this data, 500 μM and 1000 μM ITC concentrations were selected because they showed pathogen growth suppression in the case of all measured ITCs (Suppl. Table 1). Furthermore, these concentrations are known to be achievable at least transiently during biofumigation in the field (Gimsing et al., 2007; Hanschen et al., 2012; Kirkegaard & Sarwar, 1998; Matthiessen & Kirkegaard, 2006; Rudolph et al., 2015). To explore the effects of ITCs on pathogen growth alone and in combination, different ITCs were mixed in all possible two-way and three-way combinations using equal concentrations of each ITC within combinations (two-way 50:50%; three-way 33:33:33%) to achieve final low (500 μM) and high (1000 μM) ITC concentrations in 200 μl of CPG media in 96-well microplates. Microplates were cultured at 28 °C (N= 8 for all treatments) and the experiment was run for three days (72 hours), with population density measurements recorded every 24 hours as optical density at 600 nm.

### (c) Determining pathogen ITC and beta-lactam tolerance evolution in response to repeated allyl-ITC exposure

To investigate the potential for ITC tolerance evolution, we set up a 16-day selection experiment where we exposed *R. solanacearum* to 500 μM of allyl-ITC, which has the strongest effect on pathogen growth suppression of all tested ITCs (Fig. 1A; Suppl. Fig. 2). We also manipulated the frequency of ITC exposure using high (1-day), intermediate (2-day) and low (3-day) serial transfer frequency treatments. At each serial transfer, a subset of evolved bacteria (5% of the homogenised bacterial population) was serially transferred to fresh CPG media in the absence (control) and presence of allyl-ITC. ITC treatments thus manipulated both resource renewal and exposure to fresh ITC. The selection experiment was set-up following the same protocols described earlier and following this, separate fitness assays were conducted to directly compare the growth of ancestral and evolved populations (and individual colonies) in the absence and presence of 500 μM allyl-ITC. In addition to testing potential ITC tolerance evolution, we quantified changes in the growth of evolved bacteria in the absence of ITCs to reveal potential adaptations to the CPG growth media. All fitness assays were also repeated in ‘naïve’ LB media to control the potential effects of pathogen adaptation to the CPG growth media during the selection experiment. In all assays, bacteria were revived and prepared as described earlier, and grown in 96-well microplates in different media (CPG or LB) in the absence or presence of 500 μM allyl-ITC for 72 hours. Changes in ITC tolerance were quantified as bacterial growth relative to the ancestral and control treatments based on optical density at 600 nm (48-hour time point). Fitness assays were also conducted for individual bacterial colonies at the final time point where a single ancestral colony and one colony from each replicate selection line per treatment were selected resulting in a total of 49 clones.

**Figure 1.**
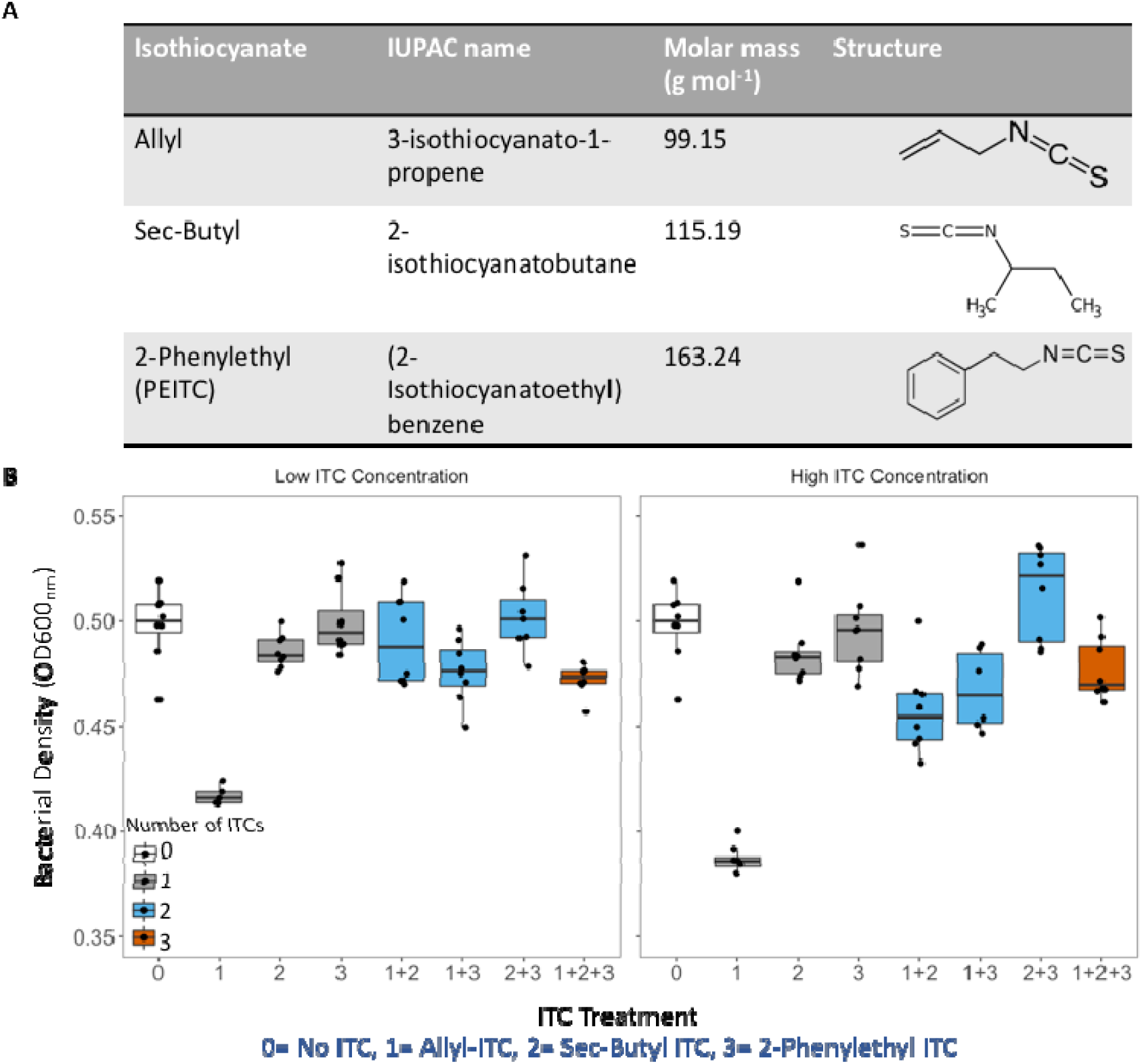
The antimicrobial activity of different ITCs against *Ralstonia solanacearum* pathogen when applied alone and in combination. The chemical properties of the three different ITCs predominantly released from Indian mustard biofumigant plant (*Brassica juncea*) (A), and their effects on *R. solanacearum* growth after 48h exposure when applied alone and in combination in liquid microcosms at low (500 μM) and high (1000 μM) concentrations (B). In (B) boxplot colours represent different ITC treatments that are labelled on X-axes as follows: (0): no-ITC (control); (1) allyl-ITC; (2) sec-butyl ITC and (3) 2- phenylethyl ITC. Individual data points show bacterial densities for each technical replicate (N=8). The boxplots show the minimum, maximum, interquartile range and the median (black line).

To explore potential ITC-tolerance mechanisms, we tested if ITC tolerance correlated with tolerance to ampicillin beta-lactam antibiotic (growth assays), which is commonly produced by various soil bacteria (Ranjan et al., 2021). Moreover, we specifically tested for ampicillin tolerance as we identified potential antibiotic-linked insertion sequence movement in our evolved clones, which has previously been shown to confer beta-lactam antibiotic tolerance in clinical settings (Boutoille et al., 2004; Poirel et al., 2003). Ampicillin tolerance was tested using the sequenced isolated clones from the final time point of the selection experiment (intermediate transfer frequency no-ITC, low transfer frequency no-ITC and low transfer frequency ITC exposure treatments) and the ancestral strain (total of 24 evolved clones and 8 replicate ancestral clones per treatment). Clones were prepared as described earlier and grown in 96-well microplates in CPG media in the absence or presence of 15 or 30 μg/ml ampicillin. Ampicillin tolerance was quantified as bacterial growth relative to the ancestral clones based on optical density at 600nm (48-hour time point).

### (d) Genome sequencing of evolved bacterial clones

A subset of evolved clones was whole genome sequenced to identify potential single nucleotide polymorphisms (SNPs), genomic rearrangements (small insertions and deletions) and potential changes in prophage and insertion sequence movement linked with *R. solanacearum* adaptation. Based on phenotypic data, we chose eight clones (1 per replicate selection line) from the low transfer frequency treatments that had evolved in the absence or presence of ITC (16 clones). Moreover, we sequenced the ancestral strain (1 clone) and eight clones from the intermediate transfer frequency no-ITC treatment (8 clones), that showed no evidence of ITC tolerance adaptation (a total of 25 clones), as controls. Genomic DNA was extracted using the Qiagen DNeasy UltraClean Microbial Kit according to the manufacturer’s protocol. DNA was quantified using the NanoDrop microvolume spectrophotometer and quality checked by gel electrophoresis imaging. DNA yields of all samples were diluted with EB buffer to 30 ng/μl concentrations and DNA samples were sent to MicrobesNG for sequencing (Illumina 30x coverage; http://www.microbesng.uk). MicrobesNG conducted library preparation using Nextera XT Library Prep Kit (Illumina, San Diego, USA) following the manufacturer’s protocol with the following modifications: 2 ng of DNA were used as input, and PCR elongation lasted 1 min. Hamilton Microlab STAR automated liquid handling system was used for DNA quantification and library preparation. Pooled libraries were quantified using the Kapa Biosystems Library Quantification Kit for Illumina on a Roche light cycler 96 qPCR machine. Libraries were sequenced on the Illumina HiSeq 2500 using a 250 bp paired end protocol. Reads were adapter trimmed using Trimmomatic 0.30 with a sliding window quality cut-off of Q15 (Bolger et al., 2014). Assembly was performed on samples using SPAdes v.3.7 (Bankevich et al., 2012) and contigs were annotated using Prokka v.1.11 (Seemann, 2014). Genomes were analysed using a standard analysis pipeline (Guarischi-Sousa et al., 2016), where reads were first mapped to a high quality and well annotated UY031 reference genome (NCBI accession: NZ_CP012687) which showed 99.95% similarity with our ancestral *R. solanacearum* strain at the chromosome level and 97.87% similarity at the mega-plasmid level. Variant calling was performed using Snippy v.3.2, a rapid haploid variant calling pipeline (Seemann, 2015). When comparing the sequenced genomes, the SNPs identified in both the ancestral strain and the evolved clones were first filtered out as these likely represent pre-existing phylogenetic differences between the reference genome and our ancestral *R. solanacearum* strain. We also compared the control treatment clones isolated from low and intermediate transfer frequency treatments (no ITC exposure) to identify potential mutations linked with CPG media adaptation. The software IMSindel v.1.0.2 (Shigemizu et al., 2018) was used to identify potential intermediate indels with options “—indelsize 10000” and using UY031 as a reference. After running IMSindel, putative indels in all isolates were combined. Putative short indels that were < 50 bp in length were removed. To investigate potential insertion sequences underlying ITC tolerance and media adaptation, insertion sequences were detected in the UY031 with ISEScan v.1.7.2.3; (Xie & Tang, 2017) using default parameters. Potential false positives were determined by blasting insertion sequences against the ISFinder database (https://isfinder.biotoul.fr/) and removing hits with an E-value > e-04. Experimental isolates were then screened for the insertion sequences identified with ISEScan using ISMapper v.2.0; (Hawkey et al., 2015) with default settings. In line with a previous study (Hawkey et al., 2020), ISMapper was run using an IS-removed UY031 assembly to improve insertion site precision. The genes flanking putative IS sites were determined by annotating the UY031 assembly using the stand-alone NCBI prokaryotic genome annotation pipeline 2021-07-01.build5508 (Tatusova et al., 2016). Additionally, we determined isolate prophage content and positions to identify potential phenotypic changes via mobile genetic elements. Isolate draft assemblies were generated using Unicycler Illumina-only assembly v.0.4.7 (Wick et al., 2017). Prophages were then identified in draft assemblies using the PHASTER (PHAge Search Tool Enhanced Release) web server (Arndt et al., 2016). Prophage movement was detected by parsing out the 5kb (or to end of contig) flanking regions either side of the prophages in the draft assemblies and mapping them to a closely related complete UY031 genome sequence. Prophage movement was detected if the flanking regions map to different parts of the UY031 genome between isolates. Prophage movement analyses were conducted using custom R and Python scripts available at (https://github.com/SamuelGreenrod/Prophage_movement). All genomes including the ancestral strain have been deposited in the European Nucleotide Archive database under the following accession number: PRJEB42551.

### (e) Statistical analysis

Repeated measures ANOVA was performed to analyse all the data with temporal sampling structure and pairwise differences were determined using *post-hoc* t-test with Bonferroni correction. All other statistical analyses (ITC tolerance and cost of tolerance in CPG and LB media and cross-tolerance in ampicillin) were conducted focusing on the 48-hour measurement time point (where ITC was still actively suppressive to *R. solanacearum*, Suppl. Fig. 1) and two-way ANOVA was used to explain variation in bacterial growth between different treatments. Tukey *post-hoc* tests were used to compare differences between subgroups (p< 0.05). Where data did not meet the assumptions of a parametric test, non-parametric Kruskal-Wallis test and post-hoc Dunn test were used. All statistical analyses and graphs were produced using R (R Foundation for Statistical Computing, R Studio v.3. 5. 1) using ggplot2, tidyverse, ggpubr, lme4, rcompanion and reshape2 packages.

## Results

### (a) Only allyl-ITC suppressed pathogen growth irrespective of the presence of other ITCs

We first determined the effects of different ITCs on *R. solanacearum* growth alone and in combination. Overall, there was a significant reduction in *R. solanacearum* densities in the presence of ITCs (ITC presence: F_1, 120_= 6.33, p< 0.01; Tukey: p< 0.05; Fig. 1B). However, this effect was mainly driven by the allyl-ITC, which significantly reduced bacterial densities compared to the no-ITC control treatment (ITC type: F_7, 114_= 49.45, p< 0.001; Tukey: p< 0.05), while other ITCs had no significant effect on the pathogen (p> 0.05; Fig. 1B). Increasing the ITC concentration from low to high (500 to 1000 μM) had no effect on inhibitory activity in either single or combination ITC treatments (ITC concentration in single ITC treatment: F_1, 43_= 2.0, p= 0.17; combination ITC treatment: F_1, 59_= 0.68, p= 0.41; Fig. 1B). However, a significant interaction between ITC type and ITC concentration in both single and combination treatments was found (ITC concentration × ITC type in single ITC treatment: F_2, 39_= 4.67, p< 0.05; in combination ITC treatment: F_3, 53_= 4.94, p< 0.01; Fig. 1B), which was driven by the increased inhibitory activity of allyl-ITC at high concentration (Tukey: p< 0.05). As a result, ITC combinations were less inhibitory than single ITC treatments (Number of ITCs: F_2, 103_= 3.82, p<0.05; Fig. 1B), which was due to reduced allyl-ITC concentration in combination treatments (total ITC concentrations were kept the same between treatments). Similarly, ITC combinations that included allyl-ITC significantly reduced bacterial densities relative to the control treatment (Allyl-ITC presence: F_1, 57_= 36.21, p< 0.001; Fig. 1B), and the presence of allyl-ITC had a clearer effect at the high ITC concentration (Allyl-ITC presence × ITC concentration: F_1, 57_= 7.51, p< 0.01; Fig. 1B). Together these results suggest that allyl-ITC was the most inhibitory compound and its antimicrobial activity was not enhanced by the presence of other ITCs.

### (b) Pathogen growth was more clearly suppressed in high and intermediate ITC exposure treatments during an experimental evolution experiment

To study the evolutionary effects of ITCs, we exposed the ancestral *R. solanacearum* strain to allyl-ITC at the low concentration (500 μM) and manipulated the frequency of exposure to ITC by transferring a subset of evolved bacterial population to fresh ITC-media mixture everyday (high), every second day (intermediate) and every third day (low) for a total of 16 days. As a result, this manipulation also affected the resource renewal rate. Overall, bacteria reached the highest population densities in the low transfer frequency treatments and the second highest in the intermediate transfer frequency treatments (Transfer frequency: F_2,45_= 4.66, p< 0.001; p< 0.05 for pairwise comparison; Fig. 2). While allyl-ITC exposure significantly reduced bacterial densities in all ITC-containing treatments (ITC presence: F_1, 46_= 30.68, p< 0.001; Fig. 2), bacterial growth was least affected in the low transfer frequency treatment (ITC presence × Transfer frequency: F_2, 42_= 4.36, p< 0.05; p< 0.001 for all pairwise comparisons; Fig. 2). The inhibitory activity of allyl-ITC also varied over time: while relatively more constant suppression was observed in the high and intermediate transfer frequency treatments, pathogen growth suppression became clear in the low transfer frequency treatment only towards the end of the selection experiment potentially due to media growth adaptation in the no-ITC control treatment (Time × Transfer frequency × ITC presence: F_2, 673_= 7.33, p< 0.001; Fig. 2). Together these results suggest that the long-term ITC activity varied temporally and depended on the ITC exposure and serial transfer frequency.

**Figure 2.**
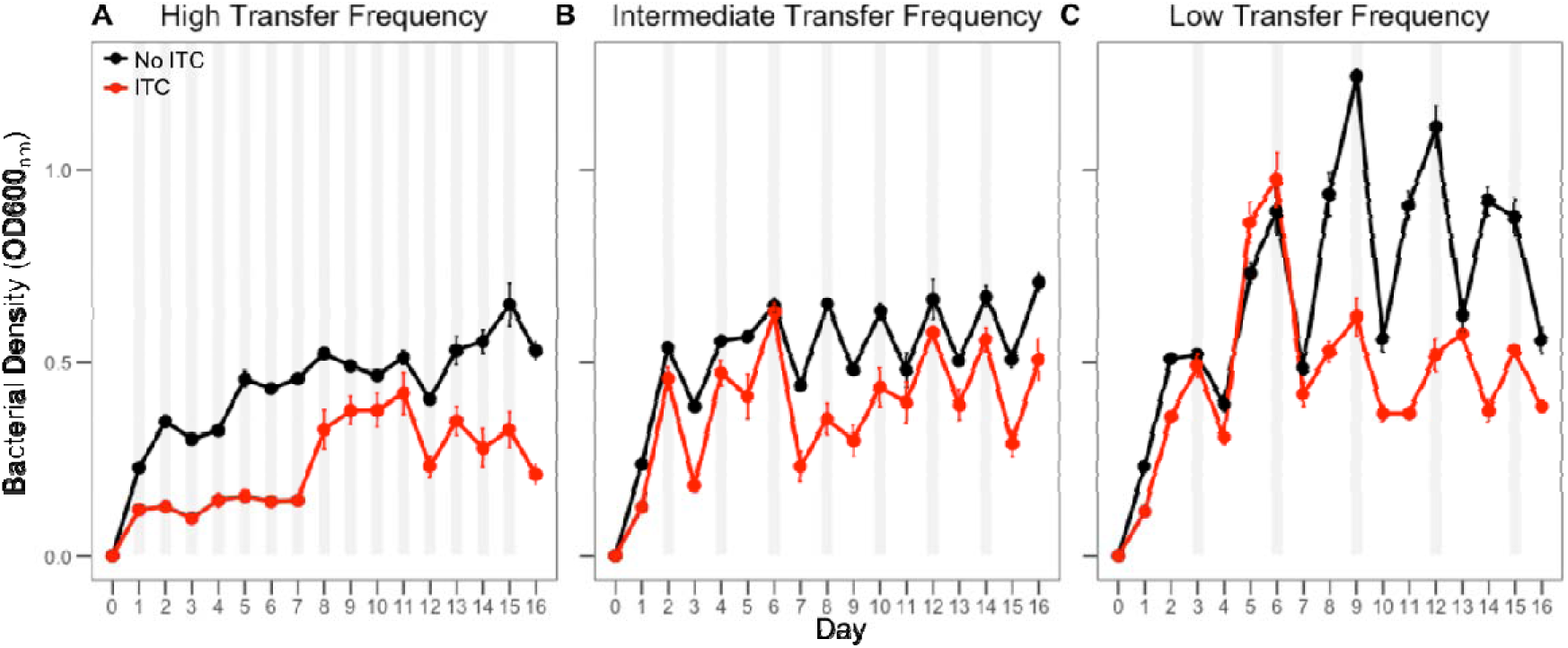
*Ralstonia solanacearum* density dynamics (OD600_nm_) during the evolution experiment in the absence and presence of allyl-ITC in high, intermediate and low transfer frequency treatments. In all panels, black and red lines correspond to *R. solanacearum* densities in the absence and presence of 500 μM allyl-ITC, respectively. Panels A-C correspond to high (1-day), intermediate (2-day) and low (3-day) transfer frequency treatments, respectively. Grey shaded areas indicate the time point of serial transfers, while optical density reads were taken at 24-hour intervals in all treatments. Each time point shows the mean of eight biological replicates and bars show ±1 error of mean.

### (c) ITC tolerance evolution was observed only in the low transfer frequency ITC exposure treatment

Fitness assays were conducted at the end of the selection experiment to compare the growth of the ancestral strain and evolved populations from different treatments in the presence and absence of allyl-ITC (experimental concentration: 500 μM). The ancestral strain reached lower densities in the presence of ITC compared to evolved populations regardless of the ITC treatment they had evolved in during the selection experiment (Evolutionary history: F_2, 45_= 5.39, p< 0.01; Tukey: p< 0.05; Fig. 3A). However, ITC tolerance was mainly observed in the low transfer frequency ITC exposure treatment, while populations that had evolved in the high or intermediate transfer frequency treatments did not significantly differ from the ancestral strain (Transfer frequency within ITC-exposed populations: F_2, 19_= 24.72, p< 0.001; Tukey: p< 0.05; Fig. 3A). Surprisingly, even the control populations that had evolved in the absence of ITCs in the low transfer frequency treatment showed an increase in ITC tolerance (p< 0.05; Fig. 3A). One potential explanation for this is that these populations adapted to grow better in CPG media, which could have helped to compensate for the mortality imposed by allyl-ITC during the fitness assays. To test this, we compared the growth of ancestral and evolved populations in the absence of allyl-ITC in the CPG media (Fig. 3B). We found that all control populations showed improved growth in the CPG media compared to ITC-exposed populations regardless of the transfer frequency treatment (Evolutionary history: F_1, 40_= 20.00, p< 0.001; Transfer frequency: F_2, 40_= 2.66, p= 0.08, in all pairwise comparisons, Tukey: p< 0.05; Fig. 3B). In contrast, none of the ITC- exposed populations showed improved growth in CPG media relative to the ancestral strain (Tukey: p< 0.05; Fig. 3B), which suggests that ITC exposure constrained *R. solanacearum* adaptation to the growth media.

**Figure 3.**
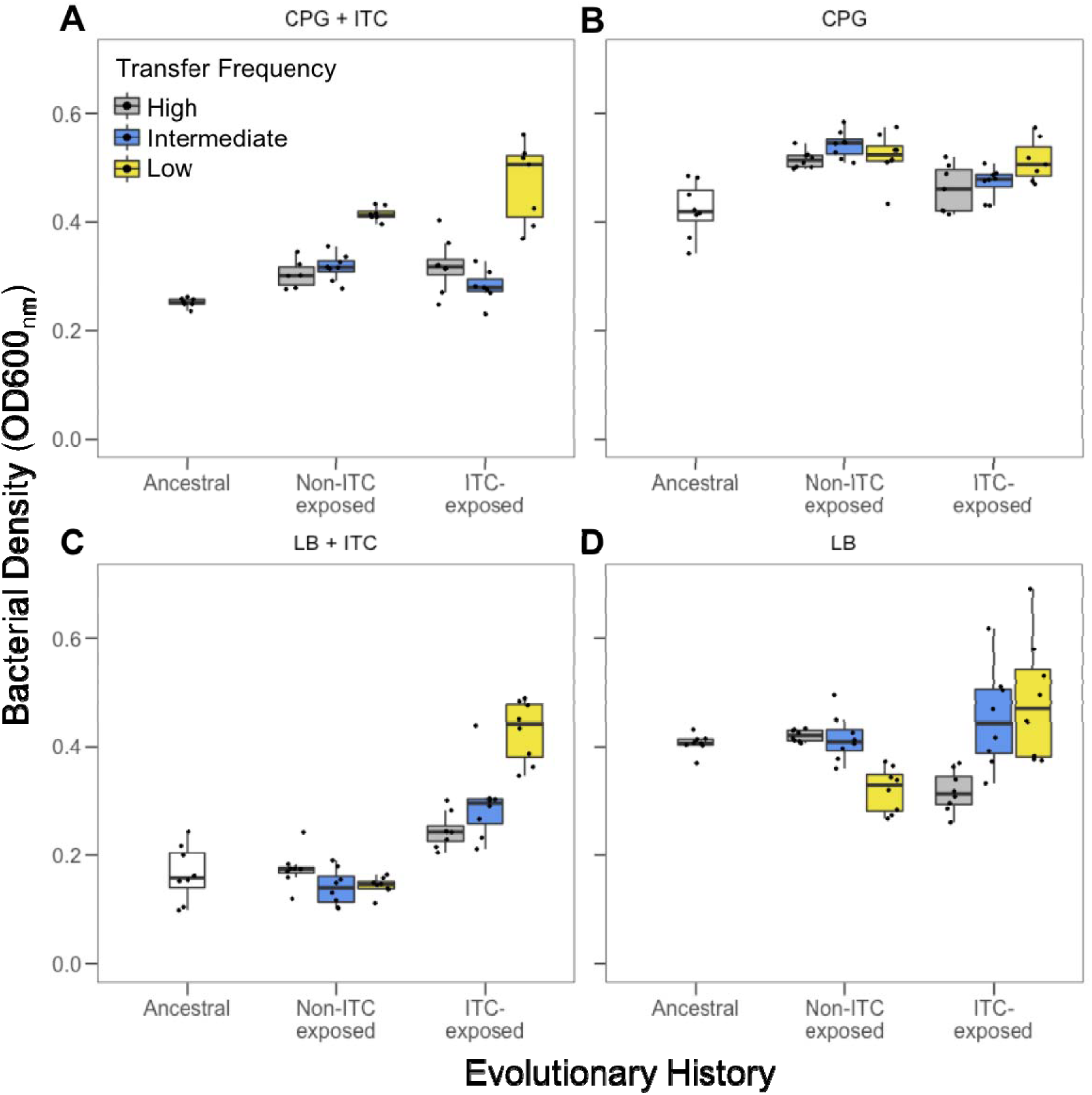
Comparison of *Ralstonia solanacearum* ITC tolerance between the ancestral clone and evolved populations from high, intermediate and low transfer frequency treatments at the end of the evolution experiment in CPG and LB media. ITC tolerance was determined as *R. solanacearum* growth (OD600_nm_) after 48h exposure to 500 μM allyl-ITC in CPG (A) and LB (C) media. Growth was also measured in the absence of allyl-ITC in both CPG (B) and LB (D) media. High (1-day), intermediate (2-day) and low (3-day) transfer frequency treatments are shown in grey, blue and yellow boxplots, respectively, and boxplots show the minimum, maximum, interquartile range and the median (black line). Individual data points show bacterial densities for each biological replicate population (N=8).

To disentangle the effects due to adaptation to the media and allyl-ITC, we repeated fitness assays in ‘naïve’ LB growth media which the bacteria had not adapted to. ITC tolerance was observed only when bacterial populations had previously been exposed to allyl-ITC (Evolutionary history: F_2, 49_= 18.82, p< 0.001; Tukey: p< 0.05; Fig. 3C), and this effect was driven by adaptation in the low transfer frequency ITC exposure treatment (no ITC tolerance was observed in the high and intermediate transfer frequency treatment; Transfer frequency: F_2, 49_= 4.37, p< 0.01; Tukey: p< 0.05; Fig. 3C). Crucially, CPG-adapted control populations showed no signs of ITC tolerance, but instead, suffered reduced growth in LB media relative to the ancestral strain and ITC-exposed populations (Evolutionary history: F_2,49_= 94.89, p< 0.001; Fig. 3D), which was clearest in the low transfer frequency exposure treatment (Evolutionary history × Transfer frequency: F_2, 49_= 23.17, p< 0.001; Fig. 3D).

We further validated our population level fitness results using individual clones (one randomly chosen clone per replicate population per treatment). In line with previous findings, ITC-exposed clones showed increased ITC tolerance compared to the control and ancestral bacterium in the LB media (Evolutionary history: F_2, 49_= 14.20, p< 0.001; Fig. 4A), and tolerance evolution was the greatest in the low transfer frequency ITC exposure treatment (Transfer frequency: F_2, 49_= 11.15, p< 0.001; Tukey: p< 0.05; Evolutionary history × Transfer frequency: F_2, 49_= 3.04, p< 0.05; Fig. 4A). Together, our results suggest that ITC tolerance, which evolved in the low transfer frequency ITC exposure treatment was robust and independent of the growth media it was quantified in. Moreover, while all control populations adapted to grow better in the CPG media, this adaptation had a positive effect on ITC tolerance only when quantified in CPG media and when the clones had evolved in the low transfer frequency treatment.

**Figure 4.**
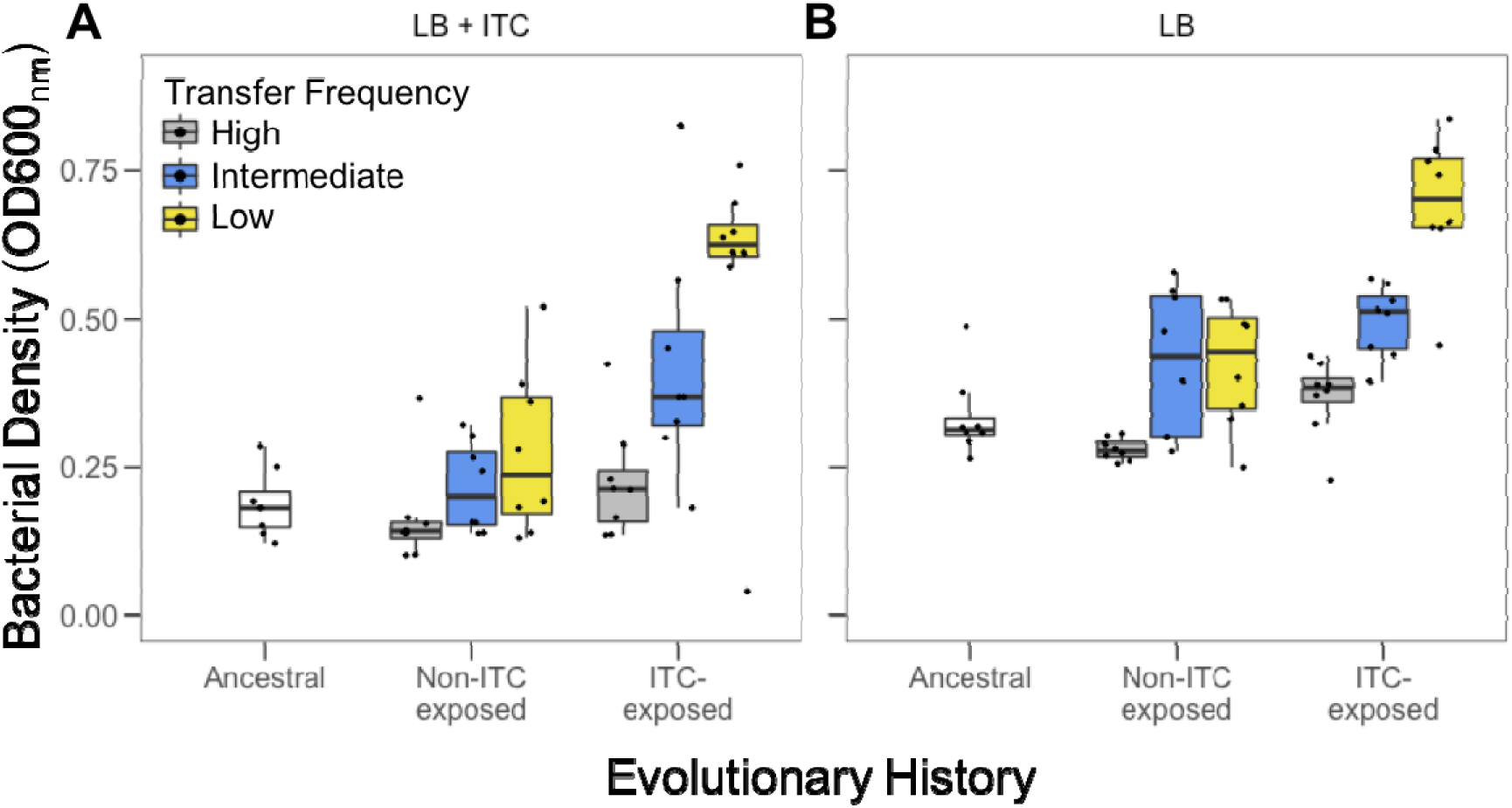
Comparison of *Ralstonia solanacearum* ITC tolerance between the ancestral and evolved clones from high, intermediate and low transfer frequency treatments at the end of the evolution experiment in LB media. ITC tolerance was determined as *R. solanacearum* growth (OD600_nm_) after 48h exposure to 500 μM allyl-ITC in LB media (A). Growth was also measured in the absence of allyl-ITC (B). High (1-day), intermediate (2-day) and low (3-day) frequency treatments are shown in grey, blue and yellow, respectively, and boxplots show the minimum, maximum, interquartile range and the median (black line). Individual data points show bacterial densities for each biological replicate population (N=8).

### (d) Evolution of ITC-tolerance confers cross-tolerance to ampicillin beta-lactam antibiotic

We also tested if exposure to allyl-ITC could have led to cross-tolerance to other antimicrobials such as the beta-lactam antibiotic ampicillin. Overall, both low (15 μg/ml) and high (30 μg/ml) ampicillin concentrations had negative effects on *R. solanacearum* growth relative to the no-ampicillin control treatment (Ampicillin concentration: F_2, 93_= 50.12, p< 0.001; Tukey: p< 0.05; high concentration was relatively more inhibitory, Suppl. Fig. 3). However, the evolved clones from the low transfer frequency ITC exposure treatment reached significantly higher bacterial densities than the ancestral strain (Evolutionary history: F_3, 92_= 3.51, p< 0.05; Tukey: p< 0.05; Suppl. Fig. 3), while evolved clones derived from low and intermediate transfer frequency control treatments (no prior ITC exposure) did not differ from the ancestral strain (Tukey: p> 0.05; Suppl. Fig. 3). Ampicillin tolerance was only observed in the high ampicillin concentration (High ampicillin concentration: F_3, 28_= 8.22, p< 0.001; Suppl. Fig. 3C; Low ampicillin concentration: F_3, 28_= 1.551, p= 0.223; Suppl. Fig. 3B). Together these results suggest that ITC tolerance conferred cross-tolerance to ampicillin for clones that had evolved in the low transfer frequency ITC exposure treatment.

### (e) Media adaptation and ITC tolerance are linked to a few mutations and loss of insertion sequences

A subset of clones which were phenotyped regarding ITC and ampicillin tolerance were selected for genome sequencing (N=25). All isolated colonies showed ancestral, fluid colony morphotype with no evidence for spontaneous evolution of small colony types as observed previously (Khokhani et al., 2017; Perrier et al., 2019). Specifically, we focused on comparing parallel small mutations and indels, intermediate indels (>50 bp) and prophage and insertion sequence (IS) movement between populations that had evolved in the absence and presence of ITC in the low transfer frequency treatments (evidence of ITC tolerance evolution) with ancestral and control populations from the intermediate transfer frequency treatment (no ITC tolerance evolution observed). Potential genetic changes were investigated in both the chromosome and megaplasmid of the bipartite genome.

Only a few mutations were observed in 1 to 6 different genes, which was expected considering the relatively short duration of the selection experiment (16 days). Of these mutations, 8 were non-synonymous and 4 synonymous (Table 1). Some mutations were observed across all treatments, indicative of adaptation to the culture media or other experimental conditions. For example, parallel non-synonymous mutations in *hisH1* gene controlling imidazole glycerol phosphate synthase were observed in 6/8 to 8/8 replicate clones in all treatments (Table 1; Fig. 5). Similarly, non-synonymous mutations in serine/threonine protein kinase genes (between 5/8 to 8/8 replicate clones) and synonymous mutations in putative deoxyribonuclease *RhsC* gene (between 1/8 to 5/8 replicate clones) were found across all treatments (Table 1; Fig. 5). A single clone that had evolved in the absence of allyl-ITC in the intermediate transfer frequency treatment had a unique non-synonymous mutation in the gene encoding the putative HTH-type transcriptional regulator *DmlR* and another clone originating from this treatment had a mutation in the IS5/IS1182 family transposase encoding gene (Table 1; Fig. 5). Additionally, we observed mutations exclusively in the low transfer control clones in genes encoding the dehydrogenase-like uncharacterised protein (3/8 replicate clones) and Tat pathway signal protein (2/8 replicate clones; Table 1; Fig. 5), which may explain ITC tolerance via media adaptation. However, no clear parallel mutations exclusive to the low frequency ITC- exposed populations were found.

**Figure 5.**
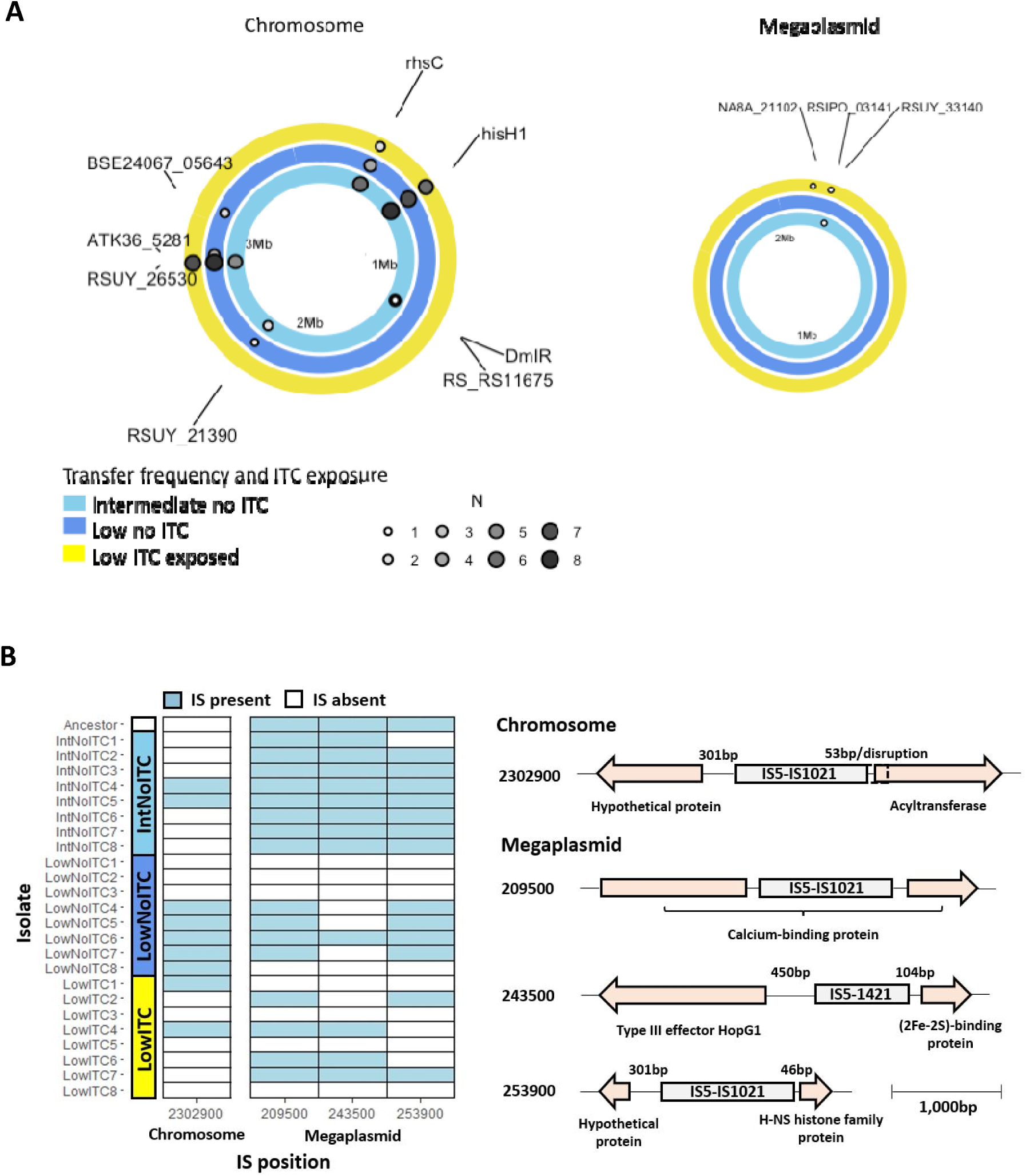
Mutations (A) and insertion sequences (IS; B) associated with evolved *Ralstonia solanacearum* clones. Each ring in panel A represents the *R. solanacearum* genome (Chromosome on the left and Megaplasmid on the right). Rings are grouped by the sequenced treatments) in different colours (see key) and dots represent mutations at different loci. Dots are sized and coloured by the number of replicates that had the same mutations (N=8) in the indicated locus. Labels show the gene name, when named, or the numbered locus tag. Distance marker is shown as Mb within each ring. In panel B, tile plot shows presence (filled tiles) and absence (unfilled tiles) of insertion sequences (ISs) in each isolate. The X-axis of the tile plot shows the IS position rounded to the nearest 100 bp. Gene schematics on the right show insertion sequence at each position and nearby genes. Gene annotation and distance between insertion sequence and genes are shown, with gene size and distance proportional to the scale bar (bottom right).

**Table 1.**
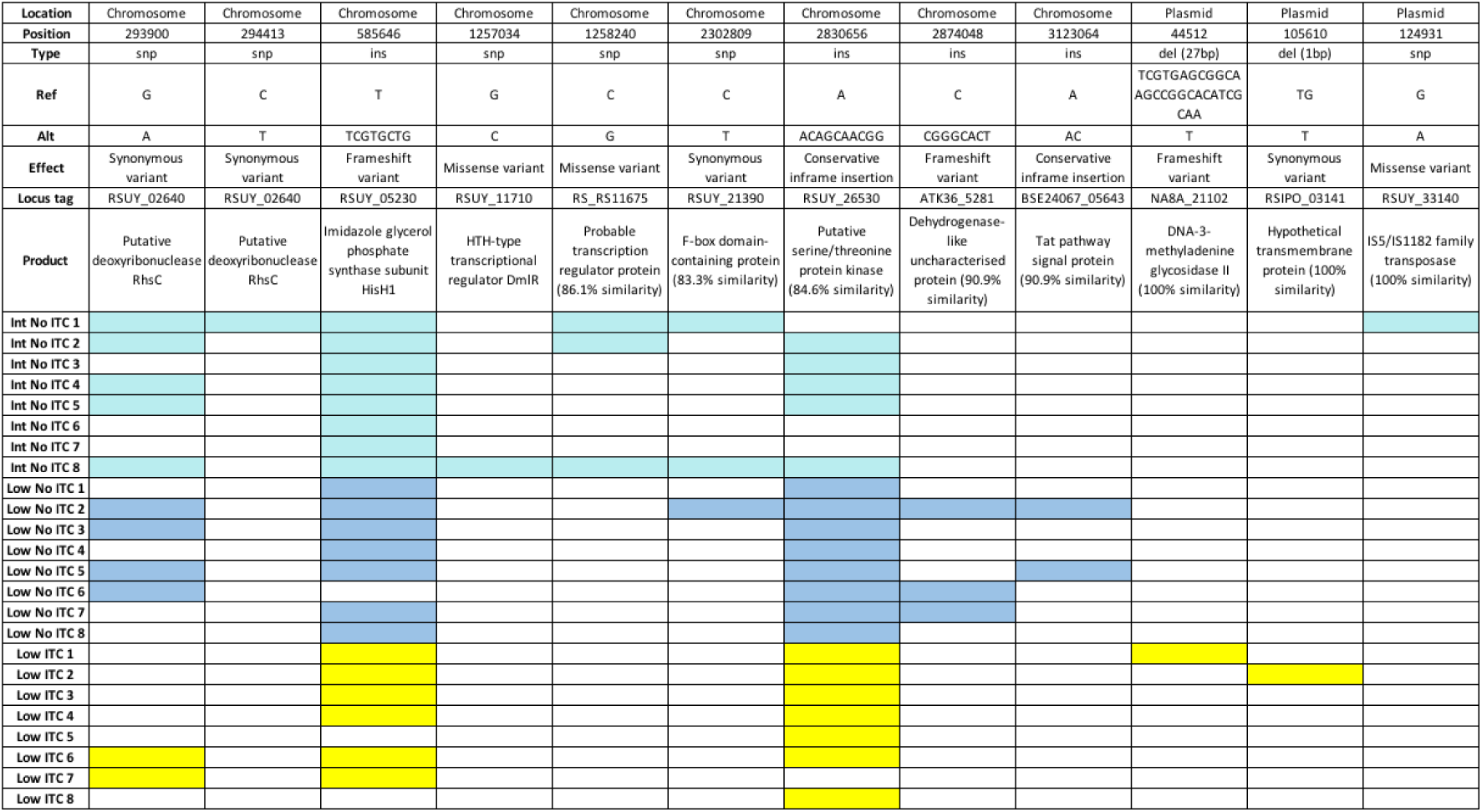
Mutated Ralstonia solanacearum genes and annotated gene functions observed in intermediate and low transfer frequency control (no-ITC), and ITC-exposed low transfer frequency treatments. Gene function predictions were derived based on BLAST using UNIPROT and percentage (%) sequence similarity is included for putative (hypothetical) proteins. Filled cells denote for the presence of mutations in given clones and white cells denote for the absence of given mutations. Replicates are named by treatments, IntNoITC= Intermediate transfer frequency, no ITC; LowNoITC= Low transfer frequency, no ITC; LowITC= Low transfer frequency, ITC.

In terms of putative intermediate indels (>50 bp), we identified 122 and 116 indel sites in the chromosome (Chr) and megaplasmid (MP), respectively. Almost all of these were insertions (Chr, 119/122; MP, 113/116) and the majority were singletons (Chr, 101/122; MP, 95/116) or doubletons (Chr, 14/122; MP, 13/116). The number of intermediate indels did not differ between evolutionary treatments either in the chromosome (Kruskal-Wallis: x^2^ = 3.65; df= 2; p= 0.161) or megaplasmid (Kruskal-Wallis: x^2^ = 3.46; df= 2; p= 0.178). As a result, this genetic variation was likely non-adaptive and driven by random drift.

To identify other potential molecular mechanisms, variation in prophages and insertion sequences (ISs) was investigated. Two prophages were found in all sequenced isolates: Inoviridae prophage φRS551 and a novel, unclassified prophage (Table S2). Prophage genome positions were almost identical between all sequenced isolates (Table S2). Therefore, no evidence for systematic prophage movement was observed in the evolved isolates relative to the ancestral strain. In contrast, ISs appeared to be highly mobile regarding 15 variable positions in the chromosome and 15 variable positions in the megaplasmid (Suppl. Fig. 5). In most variable positions (7 in the chromosome and 9 in the megaplasmid), the gain or loss of ISs was infrequent, occurring in up to three clones per treatment (Suppl. Fig. 5), which is indicative of non-adaptive, random IS movement. However, the remaining IS positions showed higher frequency of gain or loss, indicating of potentially adaptive IS movement which was also in some cases treatment-specific. For example, an IS element in position 2302900 on the chromosome absent in the ancestral strain was observed in 2 clones in the intermediate transfer frequency control and 2 low transfer ITC treatment clones, while it was gained by 5 clones in low transfer control treatment. The IS element in this position was found to be close to the start codon (∼50 bp) of an acyltransferase. In two of the low transfer control clones, the IS was found to disrupt the gene (Fig. 5), potentially knocking out acyltransferase gene expression after inserting into this position. Moreover, three IS elements in the megaplasmid were almost exclusively lost in the low transfer frequency treatment (Fig. 5). In one of the positions (209500), the IS disrupted a putative calcium-binding protein in the intermediate transfer control clones but was absent in 4/8 low transfer control and 4/8 low transfer ITC treatment clones. In the other two positions (243500 and 253900), the ISs were intergenic (positioned 450 bp and 104 bp (243500) and 301 bp and 46 bp (253900) from their left- and right-flanking genes; Fig. 5). The right-flanking genes closest to the ISs included a (2Fe-2S)-binding protein (243500) and an H-NS histone family protein (253900), whilst the left-flanking genes included the type III effector *HopG1* (243500) and an unknown hypothetical protein (253900). The frequency of IS absence in these positions (243500 and 253900) differed between low transfer treatments. Specifically, in position 243500, the IS was absent in 7/8 low transfer control and 5/8 low transfer ITC treatment clones. Meanwhile, in position 253900, the IS was absent in 4/8 low transfer control and 6/8 low transfer ITC treatment clones. However, despite these patterns, the extent of IS loss did not differ statistically between low transfer control and ITC-exposed clones when analysed individually (Mann-Whitney: 209500: w= 32, n1= 8, n2= 8, p= 1; 243500: w= 40, n1= 8, n2= 8, p= 0.29; 253900: w= 24, n1= 8, n2= 8, p= 0.35) or in combination (Mann-Whitney: w= 32, n1= 8, n2= 8, p= 1). Together, these results suggest that media adaptation and ITC tolerance was potentially driven by parallel mutations in a few genes and more frequent loss of IS elements in the low transfer frequency treatments.

## Discussion

Here we studied the effects of *Brassica*-derived ITC allelochemicals for the suppression and tolerance evolution of plant pathogenic *R. solanacearum* bacterium in a model biofumigation experiment. We found that only allyl-ITC suppressed *R. solanacearum* growth, while no reduction in pathogen densities were observed when sec-butyl and 2- phenylethyl ITCs were applied alone or in combination. By using experimental evolution, we further showed that long-term allyl-ITC exposure selected for ITC tolerance in the low transfer frequency ITC exposure treatment and was associated with cross-tolerance to ampicillin. At the genetic level, tolerance evolution was associated with the loss of IS elements. Together, our results suggest that allyl-ITC derived from Indian mustard is effective at suppressing the growth of the *R. solanacearum* pathogen *in vitro*. However, prolonged exposure could select for increased ITC tolerance, potentially reducing the efficiency of ITC-based biocontrol.

Only allyl-ITC suppressed pathogen growth and its effects were not enhanced by the presence of other ITCs. This contradicts previous studies which demonstrated *R. solanacearum* sensitivity to 2-phenylethyl ITC at concentrations as low as 330 μM (Smith & Kirkegaard, 2002). However, in the previous experiment *R. solanacearum* was exposed to 2-phenylethyl ITC in agar instead of liquid media, which has been shown to increase the toxicity of ITCs (Sarwar et al., 1998). Moreover, it is possible that different *R. solanacearum* strains respond differently to ITCs, which could also explain discrepancy between ours and other studies. While the suppressive effects of sec-butyl ITC have previously been demonstrated against dust mites (Yun et al., 2012) and fungi (Bainard et al., 2009), no antimicrobial activity has been observed in bacteria. Variation in the antimicrobial activity of ITCs could be explained by differences in chemical side-chain structure and molecular weight which govern ITC volatility and hydrophobicity (Sarwar et al., 1998). Previous studies have shown greater pathogen suppression by ITCs with aliphatic compared to aromatic sidechains in fungal pathogens (Kurt et al., 2011; Sarwar et al., 1998), insect pests (Matthiessen & Shackleton, 2005), and weeds (Vaughn et al., n.d.). With bacteria, the toxicity of allyl-ITC could be attributed to its high volatility, very short R-side chains and high reactivity (Kirkegaard & Sarwar, 1998; Manici et al., 1997; Neubauer et al., 2014). These properties could enable rapid diffusion through the liquid media before ITC is lost in the gaseous phase (Wang et al., 2009). This is supported by a study by Sarwar *et al*. (Sarwar et al., 1998), where a droplet of aliphatic allyl-ITC was shown to volatilise at room temperature in 5 minutes, whilst aromatic 2-phenylethyl ITC remained in the liquid for over 72 hours. Together, our result suggests that high volatility and reactivity could be important properties determining the antibacterial effects of ITCs.

The evolution of ITC tolerance was mainly observed in the low transfer frequency ITC exposure treatment. However, we also found that low transfer frequency control populations showed improved tolerance measured in CPG media even though they had not been exposed to allyl-ITC during the experiment. As all treatments were kept separate from each other using tightly sealed bags, this effect is unlikely explained by ‘cross selection’ due to ITC volatilisation. Alternatively, ITC tolerance evolution could have been linked to certain metabolic adaptations in this transfer frequency treatment. In support of this, we found that evolved control bacterial populations showed improved growth in the CPG media relative to ancestral and ITC-exposed populations, indicative of media adaptation. While similar media adaptations were observed in all control treatment populations, it is not clear why ITC tolerance did not evolve under one- and two-day transfer frequency treatments. One potential explanation for this could be growth-dependent effects on mutation rates. For example, prior studies have shown that bacterial mutation rates can be elevated at stationary phase (Loewe et al., 2003; Navarro Llorens et al., 2010), which could have promoted ITC tolerance and media adaptation in the low transfer frequency treatment where bacteria had spent the relatively longest time at stationary phase (Suppl. Fig. 1). Alternatively, stationary phase growth conditions could have triggered expression of stress tolerance genes, enabling selection for mutants with relatively higher ITC tolerance (Navarro Llorens et al., 2010). For example, expression of *RpoS* sigma factor in *P. aeruginosa* has previously been linked to elevated antibiotic resistance and biofilm formation at stationary phase (Murakami et al., 2005; Olsen, 2015). While more work is needed to elucidate these mechanisms, it is likely that the periodic 3-day growth cycle was important for driving ITC tolerance evolution in our experimental conditions. Interestingly, the ITC tolerance that evolved in the absence of allyl-ITC exposure was specific to CPG media and disappeared when measured in ‘naïve’ LB media. This result suggests that ITC tolerance observed in control populations was likely driven by adaptation to CPG growth media. Such adaptation may have helped to offset the suppressive effects of allyl-ITC by boosting pathogen growth to compensate increased mortality. Alternatively, it is possible that the glucose availability in the CPG media indirectly favoured the evolution of ITC tolerance via metabolic adaptations, which has previously been shown to occur both in the absence (Knöppel et al., 2017) and presence of clinical antibiotics (Zampieri et al., 2017). Together, our results suggest that prior exposure to allyl-ITC was required for the evolution of robust ITC tolerance, which was independent of the growth media.

At the genetic level, ITC tolerance was not associated with any clear parallel mutations or indels in the low transfer frequency treatments. Three clones from the low transfer frequency control treatment had unique mutations in a gene encoding a dehydrogenase-like uncharacterised protein. Dehydrogenase genes have previously been associated with both metabolism and antibiotic resistance (Marshall et al., 1999). For instance, in *Escherichia coli*, a mutation in a glucose dehydrogenase gene has been shown to function in lipopolysaccharide modification and calanic acid biosynthesis, which enabled resistance to polymyxin and other antimicrobial peptides (Lacour et al., 2008; Rodionova et al., 2020), and may have contributed to ITC tolerance in these clones. Additionally, two clones from the low transfer control treatment had mutations in a gene encoding a Tat pathway signal protein which is involved in protein translocation across membranes (Palmer et al., 2005), and may have enabled improved growth in the CPG media. Three clones from the intermediate transfer frequency treatment had unique mutations in a gene encoding a probable transcription regulator protein. While there is little information available regarding this gene, it is located beside the IS2 transposase *TnpB* gene, potentially affecting its regulation in DNA replication, recombination and repair activity (Pasternak et al., 2013). Instead of treatment-specific parallel mutations, certain mutations were found across all treatments. For example, mutations in genes encoding putative serine/threonine protein kinases, amino acid biosynthesis (*hisH1* gene) and DNA replication, recombination and repair (putative *RhsC* gene) were common for clones isolated from all treatments. Mutations observed in serine/threonine protein kinase genes could have potentially affected ITC tolerance if these enzymes were targeted by the ITCs as has been shown before in the fungus *Alternaria brassicicola* (Calmes et al., 2015), and bacterial pathogen *E. coli* (Luciano & Holley, 2009). However, as these mutations were not specific to ITC-treatment clones, they were probably associated with bacterial growth and metabolism.

In *R. solanacearum*, insertion sequences (ISs) have been shown to affect host virulence and phenotypic plasticity by inserting into and disrupting type III effectors and global virulence regulators (Gonçalves et al., 2020; Jeong & Timmis, 2000). Therefore, we investigated whether IS movement may be the cause of *R. solanacearum* ITC tolerance adaptation. We identified one IS position in the chromosome and three positions in the megaplasmid which showed treatment specific patterns. The gain of IS at position 2302900 was primarily observed with low transfer control isolates and was situated either ∼50 bp from the start codon or inside of a putative acyltransferase. Acyltransferases have a broad range of functions including lipid storage (Ohlrogge & Browse, 1995), phospholipid biosynthesis (Li et al., 2017), and the production of toxins (Greene et al., 2015) and antibiotics (Kozakai et al., 2020). Whilst many of these functions are critical to cell growth, some such as the production of toxins would be redundant when grown in media. Therefore, gene disruption by ISs in the low transfer control may increase fitness by allowing energy and nutrients to be re-directed towards promoting cell growth and competitivity, potentially at the expense of reduced virulence *in planta*. We also found loss of two ISs in the intergenic region of the megaplasmid in the low transfer control and ITC treatments. While these were intergenic, they were close (∼50-100 bp) to the start codons of their right flanking genes and could have affected gene expression. In position 243500, the IS was situated close to a (2Fe-2S)-binding protein gene. Iron-sulfur clusters have been implicated in cellular metabolism, protein structural stabilisation, iron storage, and the regulation of gene expression (Johnson et al., 2005). In the other position (253900), the IS was situated close to an H-NS histone like protein gene and while non-significant, was lost more frequently across low transfer ITC treatment clones (6/8) than low transfer control isolates (4/8). H-NS histone like proteins are transcriptional repressors generally involved in adaptation to environmental challenges like temperature stress and osmolarity gradients (Atlung & Ingmer, 1997). Further, H-NS histone like proteins have been shown to stabilise the sigma factor *RpoS* (Hommais et al., 2001) which acts as a master regulator of the bacterial stress response. Whilst the H-NS histone-like protein could affect ITC tolerance by mediating the bacterial stress response, the impact of the (2Fe-2S)-binding protein is less clear. Notably, in *Campylobacter jejuni*, genes containing iron-sulfur clusters have been found to be upregulated in response to ITCs, potentially due to their susceptibility to oxidative stress caused by ITC exposure (Dufour et al., 2013). Therefore, by altering the expression of the (2Fe-2S)-binding protein, IS loss could increase the pool of cellular iron-sulfur cluster proteins and compensate for losses caused by ITC oxidative stress. In the final megaplasmid IS position (209500), we identified a loss of IS from a calcium-binding protein gene, which had likely disrupted gene expression or protein function in this gene with the ancestral strain. In human breast cancer cells, ITCs, including phenethyl- (Tusskorn et al., 2013) and allyl-ITC (Bo et al., 2016) have been found to induce mitochondrial calcium ion mobilisation resulting in cytotoxicity through a reduction in mitochondrial membrane potential. Whilst further work is required to determine the causes of ITC cytotoxicity in *R. solanacearum*, upregulation of calcium-binding protein gene expression could have increased ITC tolerance by facilitating the sequestration of free calcium ions. However, like other genetic changes, loss of this IS did not occur statistically more often in the presence of ITC selection. As a result, specific genetic mechanisms underlying ITC tolerance remain elusive.

In conclusion, our findings demonstrate that allyl-ITC could potentially be used to suppress the growth of *R. solanacearum* plant pathogen. However, repeated ITC exposure could select for mutants with increased ITC tolerance, potentially weakening the long-term efficiency of ITCs and biofumigation. Future work should focus on validating these findings in more complex natural environments. For example, it is currently not clear if *R. solanacearum* ITC tolerance evolves in the plant rhizosphere in the presence of other microbes that could constrain mutation supply rate via resource and direct competition. Moreover, different resistance mechanisms could be selected depending on soil physiochemical properties and nutrient and plant root exudate availability, while it is not clear if the ITC concentrations used in this experiment are achievable through biofumigation and whether they might have negative effects on beneficial soil microbes. More efficient ITC application could be attained by drilling the biofumigant plants into fields at the time of flowering when GSL levels are highest using finely chopped plant material, which maximises cell disruption and ITC release to the soil (Back et al., 2019). In addition, the efficacy of *Brassica*-based biofumigation could potentially be improved by using plant cultivars with elevated levels of sinigrin, the GSL precursor to allyl-ITC. Comprehensive *in vivo* work is thus required to validate the potential of allyl-ITC for *R. solanacearum* biocontrol in the field. It would also be interesting to study if ITC tolerance leads to life-history traits in *R. solanacearum*, potentially affecting its virulence or competitiveness in the rhizosphere.

## Acknowledgements

This work was funded by the NERC ACCE DTP (CLA), the Royal Society (RSG\R1\180213 and CHL\R1\180031; V-P.F) and jointly by a grant from UKRI, Defra, and the Scottish Government, under the Strategic Priorities Fund Plant Bacterial Diseases programme (BB/T010606/1; V-P.F) at the University of York. The funders had no role in study design, data collection and interpretation, or the decision to submit the work for publication.

## Supplementary Materials

**Supplementary Table 1.** The mean density reduction (%) of *Ralstonia solanacearum* bacterium when exposed to 500 or 1000 µM allyl, sec-butyl and 2-phenylethyl ITCs in CPG growth media after 24, 48 or 72 hours relative to when grown in the absence of ITCs. This table is based on the same data presented in Supplementary Fig. 2.

| ITC Type and Concentration ($\mu\text{M}$) | Time (h) | Bacterial density reduction (%) compared to control |
| --- | --- | --- |
| Allyl-ITC, 500 | 24 | 66 |
|  | 48 | 54 |
|  | 72 | 27 |
| Allyl-ITC, 1000 | 24 | 66 |
|  | 48 | 47 |
|  | 72 | 41 |
| Sec-Butyl ITC, 500 | 24 | 33 |
|  | 48 | 26 |
|  | 72 | 9 |
| Sec-Butyl ITC, 1000 | 24 | 30 |
|  | 48 | 27 |
|  | 72 | 8 |
| 2-Phenylethyl ITC, 500 | 24 | 39 |
|  | 48 | 13 |
|  | 72 | 10 |
| 2-Phenylethyl ITC, 1000 | 24 | 38 |
|  | 48 | 18 |
|  | 72 | 13 |

**Supplementary Table 2.** Prophage information of ancestral and experimental isolate assemblies as determined using flanking regions mapped to UY031. Replicates are named by treatments, IntNoITC= Intermediate transfer frequency, no ITC; LowNoITC= Low transfer frequency, no ITC; LowITC= Low transfer frequency, ITC.

| Clone | Prophage | Left flank UY031 position | Right flank UY031 position | Length (kb) | GC content (%) | Total proteins # |
| --- | --- | --- | --- | --- | --- | --- |
| UY031 | Unclassified A | - | - | 37 | 62.76 | 44 |
|  | RS551 | - | - | 13.4 | 61.24 | 16 |
|  | PHAGE_Vibrio_VHML_NC_004456 | - | - | 18.5 | 64.64 | 29 |
| Ancestor | Unclassified A | NZ_CP012687.1:1121824-1126823 | NZ_CP012687.1:1162291-1162775 | 35.4 | 62.85 | 42 |
|  | RS551 | NZ_CP012687.1:1218220-1223219 | NZ_CP012687.1:1236384-1237639 | 13.1 | 58.58 | 17 |
| IntNoITC1 | Unclassified A | NZ_CP012687.1:1121824-1126823 | NZ_CP012687.1:1162291-1162775 | 35.4 | 62.85 | 42 |
|  | RS551 | NZ_CP012687.1:1218233-1223232 | NZ_CP012687.1:1236385-1237640 | 13.1 | 58.58 | 17 |
| IntNoITC2 | Unclassified A | NZ_CP012687.1:1121824-1126823 | NZ_CP012687.1:1162291-1162775 | 35.4 | 62.85 | 42 |
|  | RS551 | NZ_CP012687.1:1218233-1223232 | NZ_CP012687.1:1236385-1237640 | 13.1 | 58.58 | 17 |
| IntNoITC3 | Unclassified A | NZ_CP012687.1:1121823-1126822 | NZ_CP012687.1:1162290-1162775 | 35.4 | 62.85 | 43 |
|  | RS551 | NZ_CP012687.1:1218233-1223232 | NZ_CP012687.1:1236385-1237640 | 13.1 | 58.58 | 17 |
| IntNoITC4 | Unclassified A | NZ_CP012687.1:1121823-1126822 | NZ_CP012687.1:1162290-1162775 | 35.4 | 62.85 | 43 |
|  | RS551 | NZ_CP012687.1:1218233-1223232 | NZ_CP012687.1:1236385-1237640 | 13.1 | 58.58 | 18 |
| IntNoITC5 | Unclassified A | NZ_CP012687.1:1121824-1126823 | NZ_CP012687.1:1162291-1162775 | 35.4 | 62.85 | 42 |
|  | RS551 | NZ_CP012687.1:1218233-1223232 | NZ_CP012687.1:1236385-1237640 | 13.1 | 58.58 | 18 |
| IntNoITC6 | Unclassified A | NZ_CP012687.1:1121823-1126822 | NZ_CP012687.1:1162290-1162775 | 35.4 | 62.85 | 43 |
|  | RS551 | NZ_CP012687.1:1218233-1223232 | NZ_CP012687.1:1236385-1237640 | 13.1 | 58.58 | 18 |
| IntNoITC7 | Unclassified A | NZ_CP012687.1:1121824-1126823 | NZ_CP012687.1:1162291-1162775 | 35.4 | 62.85 | 43 |
|  | RS551 | NZ_CP012687.1:1218233-1223232 | NZ_CP012687.1:1218233-1223232 | 13.1 | 58.58 | 17 |
| IntNoITC8 | Unclassified A | NZ_CP012687.1:1121823-1126822 | NZ_CP012687.1:1162290-1162775 | 35.4 | 62.85 | 41 |
|  | RS551 | NZ_CP012687.1:1218233-1223232 | NZ_CP012687.1:1236385-1237640 | 13.1 | 58.58 | 17 |
| LowNoITC1 | Unclassified A | NZ_CP012687.1:1121824-1126823 | NZ_CP012687.1:1162291-1162775 | 35.4 | 62.85 | 42 |
|  | RS551 | NZ_CP012687.1:1218233-1223232 | NZ_CP012687.1:1236385-1237640 | 13.1 | 58.58 | 17 |
| LowNoITC2 | Unclassified A | NZ_CP012687.1:1121824-1126823 | NZ_CP012687.1:1162291-1162775 | 35.4 | 62.85 | 42 |
|  | RS551 | NZ_CP012687.1:1218233-1223232 | NZ_CP012687.1:1236385-1237640 | 13.1 | 58.58 | 18 |
| LowNoITC3 | Unclassified A | NZ_CP012687.1:1121823-1126822 | NZ_CP012687.1:1162290-1162775 | 35.4 | 62.85 | 41 |
|  | RS551 | NZ_CP012687.1:1218233-1223232 | NZ_CP012687.1:1236385-1237640 | 13.1 | 58.58 | 18 |
| LowNoITC4 | Unclassified A | NZ_CP012687.1:1121823-1126822 | NZ_CP012687.1:1162290-1162775 | 35.4 | 62.85 | 43 |
|  | RS551 | NZ_CP012687.1:1218233-1223232 | NZ_CP012687.1:1236385-1237640 | 13.1 | 58.58 | 17 |
| LowNoITC5 | Unclassified A | NZ_CP012687.1:1121823-1126822 | NZ_CP012687.1:1162290-1162775 | 35.4 | 62.85 | 43 |
|  | RS551 | NZ_CP012687.1:1218220-1223219 | NZ_CP012687.1:1236385-1237640 | 13.1 | 58.58 | 18 |
| LowNoITC6 | Unclassified A | NZ_CP012687.1:1121823-1126822 | NZ_CP012687.1:1162290-1162775 | 35.4 | 62.85 | 41 |
|  | RS551 | NZ_CP012687.1:1218220-1223219 | NZ_CP012687.1:1236384-1237640 | 13.1 | 58.58 | 18 |
| LowNoITC7 | Unclassified A | NZ_CP012687.1:1121823-1126822 | NZ_CP012687.1:1162290-1162775 | 35.4 | 62.85 | 41 |
|  | RS551 | NZ_CP012687.1:1218220-1223219 | NZ_CP012687.1:1236384-1237640 | 13.1 | 58.58 | 18 |
| LowNoITC8 | Unclassified A | NZ_CP012687.1:1121824-1126823 | NZ_CP012687.1:1162291-1162775 | 35.4 | 62.85 | 42 |
|  | RS551 | NZ_CP012687.1:1218220-1223219 | NZ_CP012687.1:1236384-1237640 | 13.1 | 58.58 | 17 |
| LowITC1 | Unclassified A | NZ_CP012687.1:1121824-1126823 | NZ_CP012687.1:1162291-1162775 | 35.4 | 62.85 | 43 |
|  | RS551 | NZ_CP012687.1:1218220-1223219 | NZ_CP012687.1:1236384-1237640 | 13.1 | 58.58 | 17 |
| LowITC2 | Unclassified A | NZ_CP012687.1:1121823-1126822 | NZ_CP012687.1:1162290-1162775 | 35.4 | 62.85 | 43 |
|  | RS551 | NZ_CP012687.1:1218220-1223219 | NZ_CP012687.1:1236384-1237640 | 13.1 | 58.58 | 17 |
| LowITC3 | Unclassified A | NZ_CP012687.1:1121824-1126823 | NZ_CP012687.1:1162291-1162775 | 35.4 | 62.85 | 42 |
|  | RS551 | NZ_CP012687.1:1218220-1223219 | NZ_CP012687.1:1236384-1237640 | 13.1 | 58.58 | 18 |
| LowITC4 | Unclassified A | NZ_CP012687.1:1121824-1126823 | NZ_CP012687.1:1162291-1162775 | 35.4 | 62.85 | 42 |
|  | RS551 | NZ_CP012687.1:1218220-1223219 | NZ_CP012687.1:1236384-1237639 | 13.1 | 58.58 | 17 |
| LowITC5 | Unclassified A | NZ_CP012687.1:1121823-1126822 | NZ_CP012687.1:1162290-1162775 | 35.4 | 62.85 | 41 |
|  | RS551 | NZ_CP012687.1:1218220-1223219 | NZ_CP012687.1:1236384-1237639 | 13.1 | 58.58 | 18 |
| LowITC6 | Unclassified A | NZ_CP012687.1:1121824-1126823 | NZ_CP012687.1:1162291-1162775 | 35.4 | 62.85 | 42 |
|  | RS551 | NZ_CP012687.1:1218220-1223219 | NZ_CP012687.1:1236384-1237639 | 13.1 | 58.58 | 17 |
| LowITC7 | Unclassified A | NZ_CP012687.1:1121823-1126822 | NZ_CP012687.1:1162290-1162775 | 35.4 | 62.85 | 41 |
|  | RS551 | NZ_CP012687.1:1218220-1223219 | NZ_CP012687.1:1236384-1237639 | 13.1 | 58.58 | 18 |
| LowITC8 | Unclassified A | NZ_CP012687.1:1121823-1126822 | NZ_CP012687.1:1162290-1162775 | 35.4 | 62.85 | 41 |
|  | RS551 | NZ_CP012687.1:1218220-1223219 | NZ_CP012687.1:1236384-1237639 | 13.1 | 58.58 | 18 |

**Supplementary Figure 1.**
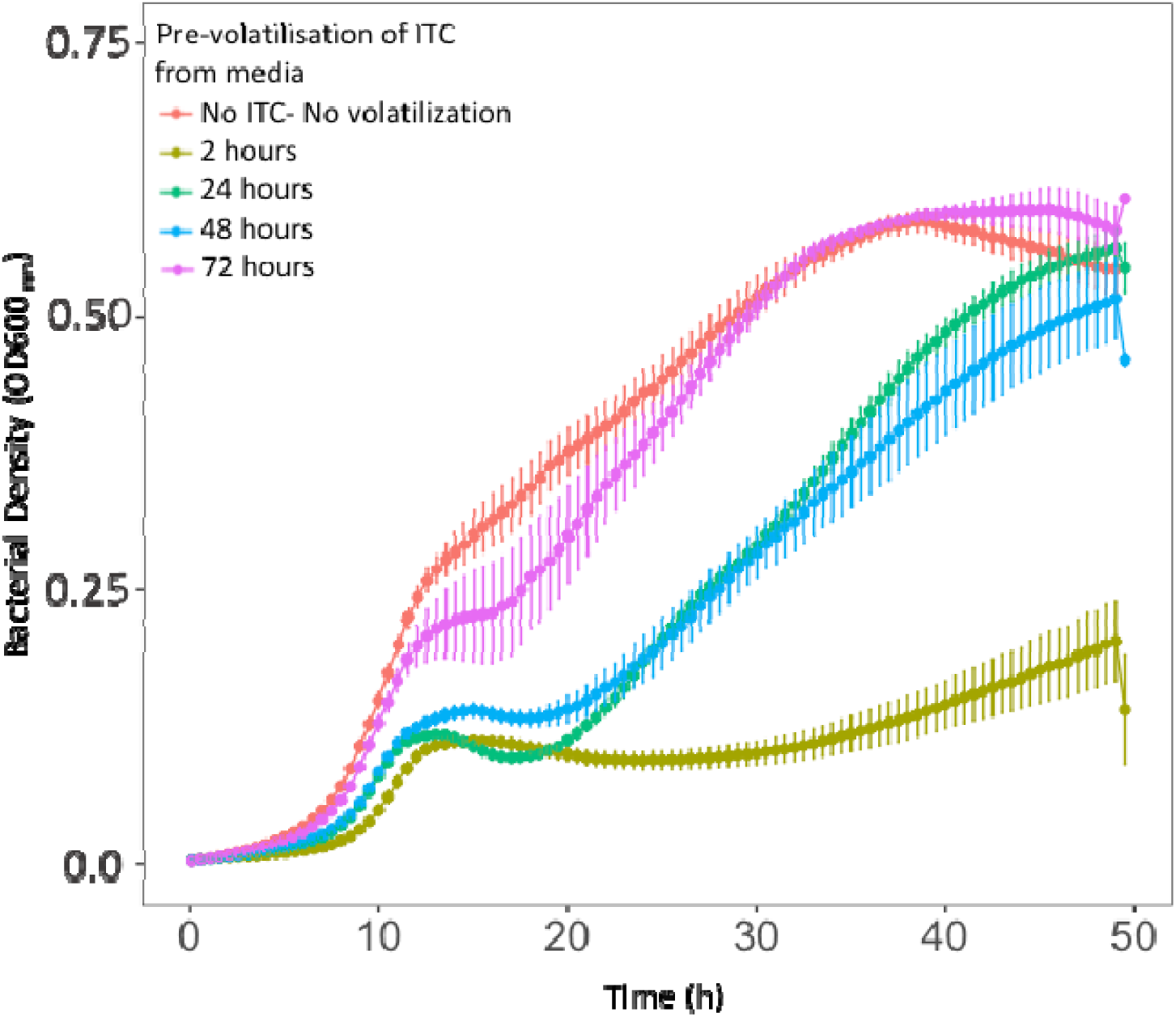
The effect of allyl-ITC pre-volatilisation for antibacterial activity against *Ralstonia solanacearum*. *R. solanacearum* bacterial growth was measured in CPG media supplemented with 0 (No allyl-ITC) or 500 μM of allyl-ITC that had been allowed to volatilise for 2, 24, 48 or 72 hours (see key). All data points show the mean of eight technical replicates and bars show ±1 standard error of the mean (SEM).

**Supplementary Figure 2.**
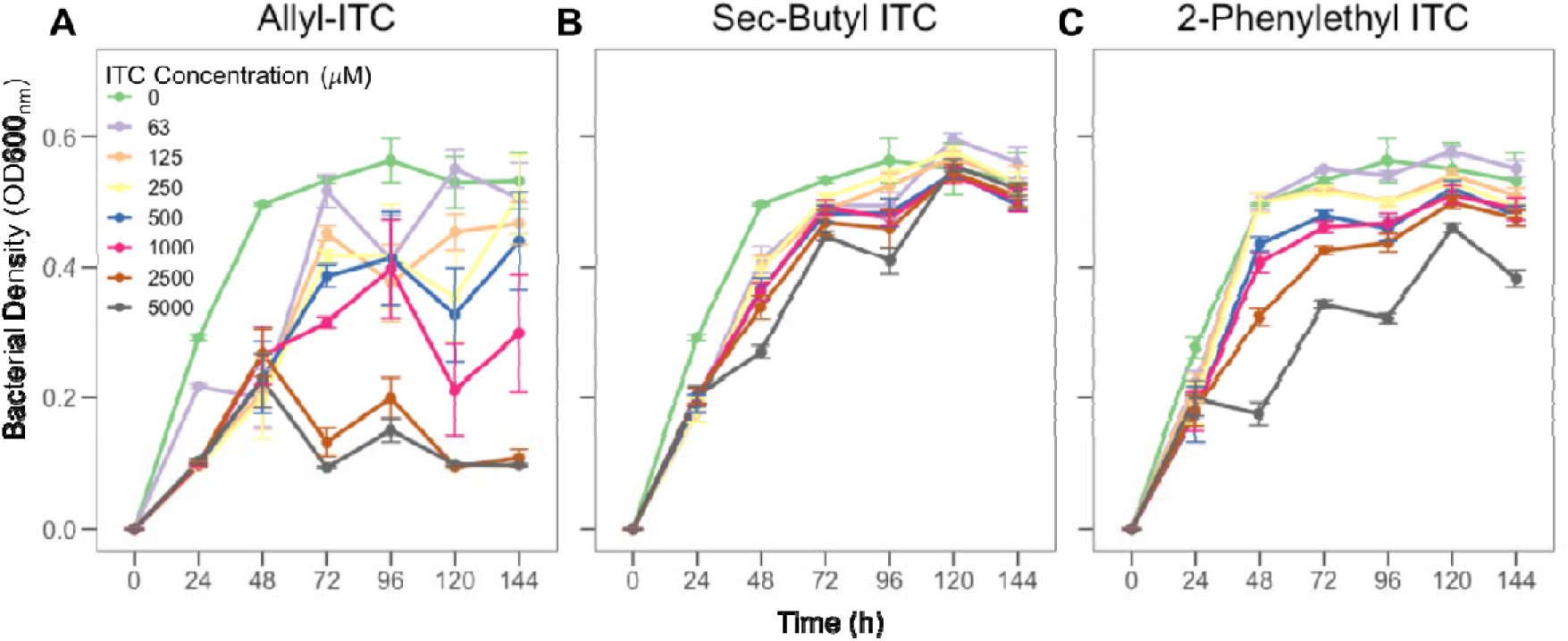
Effects of allyl, sec-butyl and 2-phenylethyl ITCs on *Ralstonia solanacearum* growth at different ITC concentrations. In all panels, *R. solanacearum* bacterial densities are shown on the Y-axis as optical density (OD600_nm_), measured at 24- hour intervals (X-axis). In all panels, different line colours refer to different ITC concentrations (see key in A). All data points show the mean of eight technical replicates and bars show ±1 standard error of the mean (SEM).

**Supplementary Figure 3.**
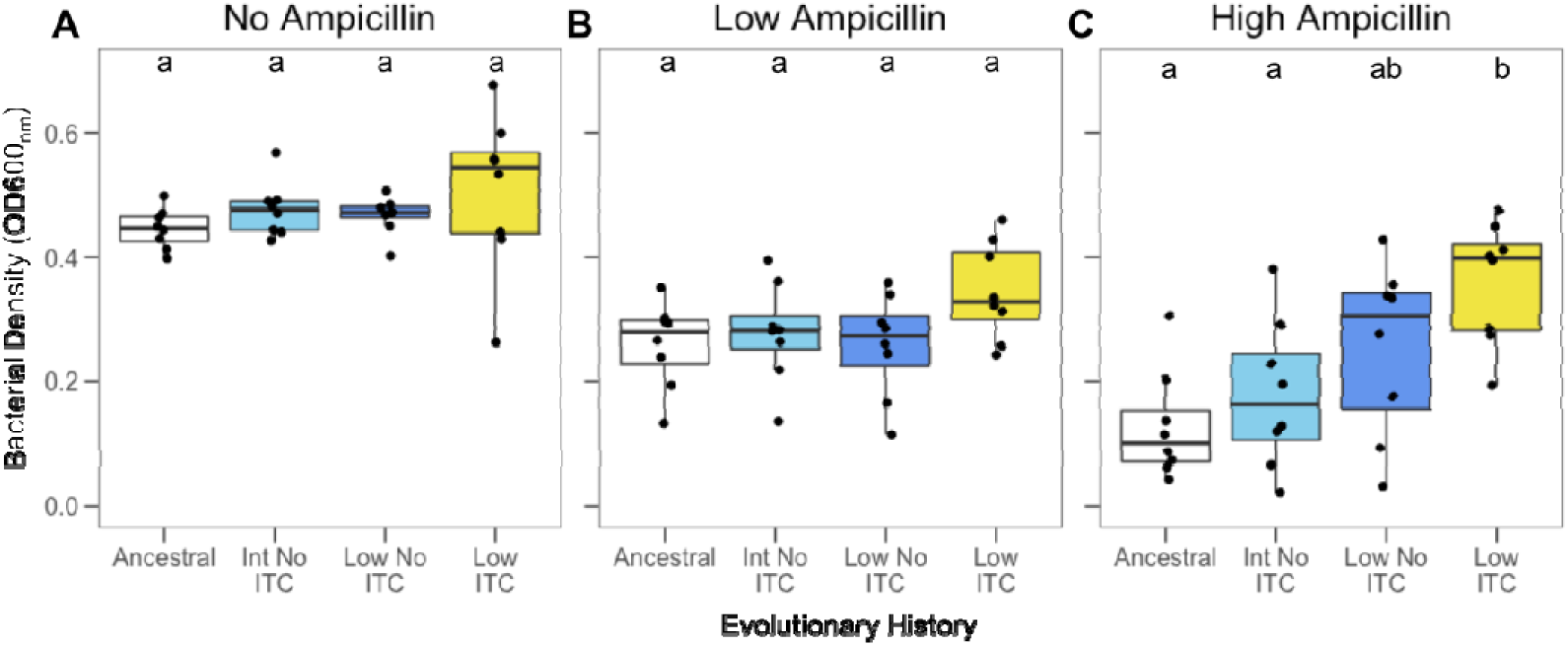
*Ralstonia solanacearum* tolerance to ampicillin beta-lactam antibiotic. Ampicillin tolerance was measured as the growth of ancestral and evolved *R. solanacearum* clones isolated from intermediate (Int) and low transfer frequency (Low) control treatments (no-ITC) and ITC-exposed low transfer frequency treatment in the absence (A) and presence (B-C) of ampicillin (15 and 30 µg/ml concentrations). Boxplots show the minimum, maximum, interquartile range and the median (black line) after 48 hours. Individual data points show bacterial densities for each biological replicate clone (N= 8). Different small case letters above boxplots indicate significant pairwise differences (Tukey: p< 0.05) between treatments within each panel.

**Supplementary Figure 4.**
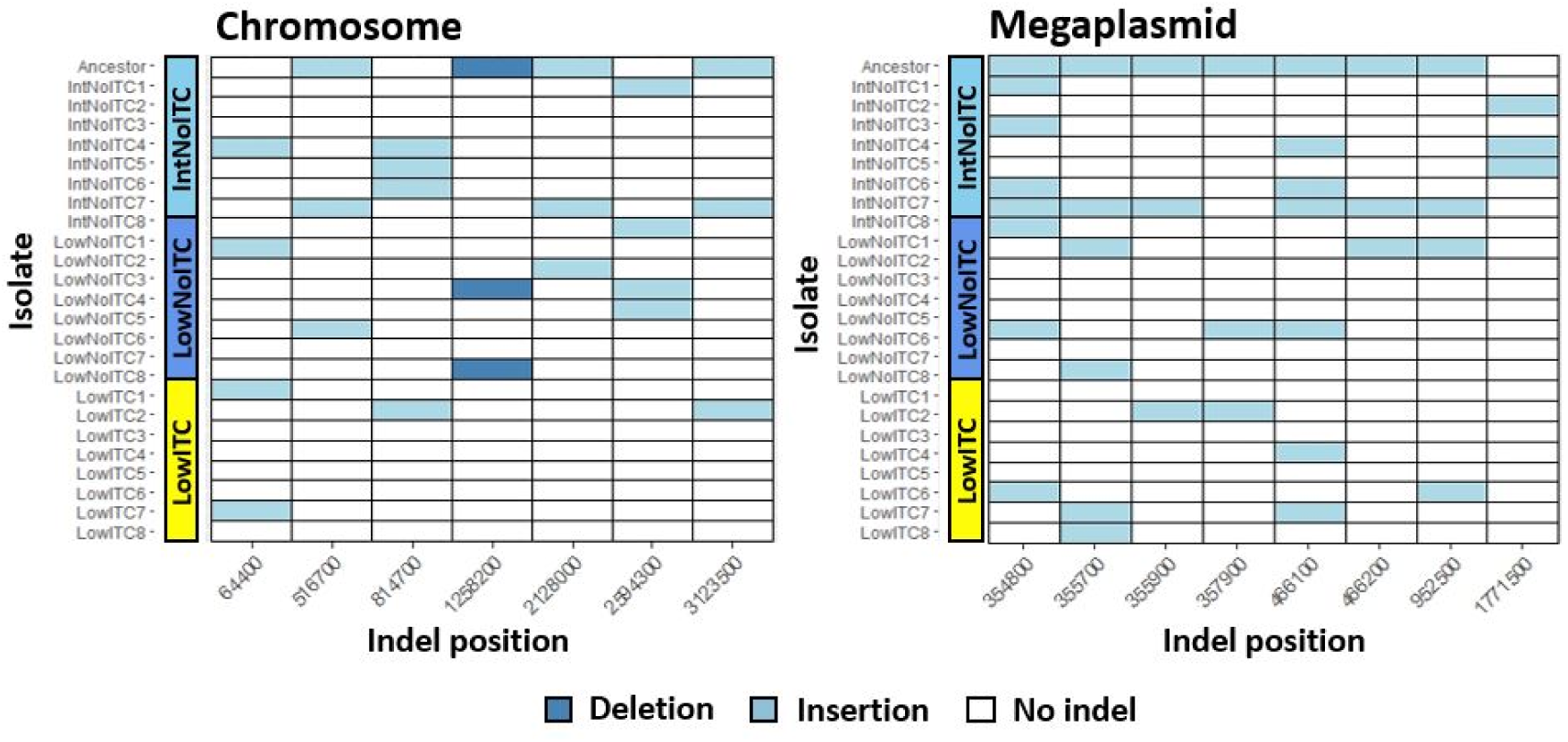
Presence and absence of intermediate indels found in more than two isolates in the chromosome and megaplasmid. The X-axis shows the indel position rounded to the nearest 100bp. The Y-axis shows isolates grouped as shown in Figure 5.

**Supplementary Figure 5.**
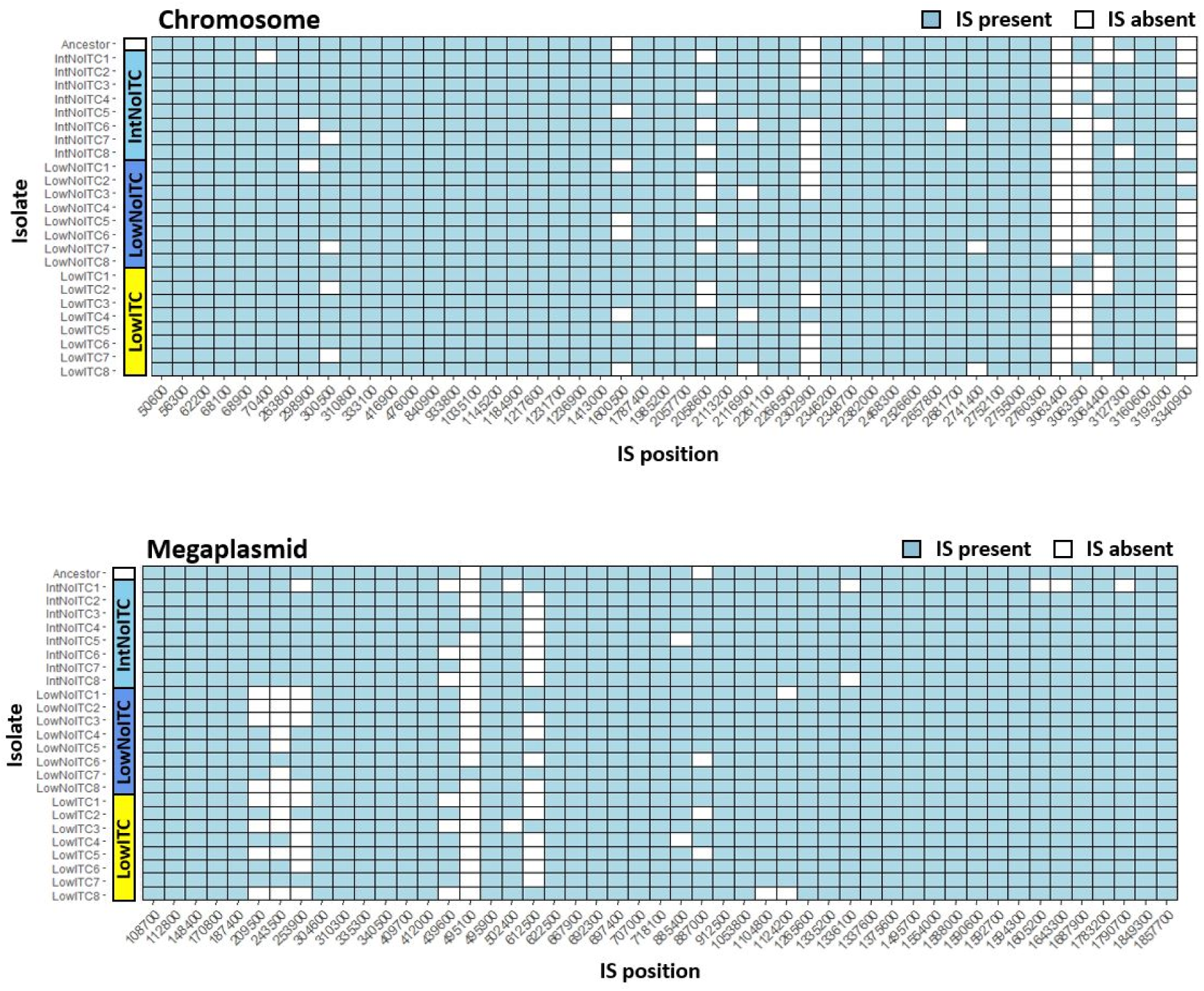
Presence and absence of insertion sequences in the chromosome and megaplasmid. The X and Y-axes show the insertion sequence position and experimental isolate, respectively, as outlined in Figure 5.

## Notes

### Competing Interest Statement

The authors have declared no competing interest.

## References

Angus, J. F., Gardner, P. A., Kirkegaard, J. A., & Desmarchelier, J. M. (1994). Biofumigation: Isothiocyanates released frombrassica roots inhibit growth of the take-all fungus. Plant and Soil, 162(1), 107–112. https://doi.org/10.1007/BF01416095

Arndt, D., Grant, J. R., Marcu, A., Sajed, T., Pon, A., Liang, Y., & Wishart, D. S. (2016). PHASTER: a better, faster version of the PHAST phage search tool. Nucleic Acids Research, 44(W1), W16–W21. https://doi.org/10.1093/nar/gkw387

Atlung, T., & Ingmer, H. (1997). H-NS: a modulator of environmentally regulated gene expression. Molecular Microbiology, 24(1), 7–17. https://doi.org/10.1046/J.1365-2958.1997.3151679.X

Back, M., Barker, A., & Evans, K. (2019). *Effectiveness of biofumigant crops for the management of PCN in GB*. Agricultural And Horticultural Development Board (AHDB). https://pure.sruc.ac.uk/en/publications/effectiveness-of-biofumigant-crops-for-the-management-of-pcn-in-g

Bainard, L. D., Brown, P. D., & Upadhyaya, M. K. (2009). Inhibitory Effect of Tall Hedge Mustard ( *Sisymbrium loeselii* ) Allelochemicals on Rangeland Plants and Arbuscular Mycorrhizal Fungi. Weed Science, 57(4), 386–393. https://doi.org/10.1614/WS-08-151.1

Bangarwa, S. K., Norsworthy, J. K., Mattice, J. D., & Gbur, E. E. (2011). Glucosinolate and Isothiocyanate Production from Brassicaceae Cover Crops in a Plasticulture Production System. Weed Science, 59(2), 247–254. https://doi.org/10.1614/WS-D-10-00137.1

Bankevich, A., Nurk, S., Antipov, D., Gurevich, A. A., Dvorkin, M., Kulikov, A. S., Lesin, V. M., Nikolenko, S. I., Pham, S., Prjibelski, A. D., Pyshkin, A. V., Sirotkin, A. V., Vyahhi, N., Tesler, G., Alekseyev, M. A., & Pevzner, P. A. (2012). SPAdes: A new genome assembly algorithm and its applications to single-cell sequencing. Journal of Computational Biology, 19(5), 455–477. https://doi.org/10.1089/cmb.2012.0021

Bending, G. D., & Lincoln, S. D. (1999). Characterisation of volatile sulphur-containing compounds produced during decomposition of Brassica juncea tissues in soil. Soil Biology and Biochemistry, 31(5), 695–703. https://doi.org/10.1016/S0038-0717(98)00163-1

Bo, P., Lien, J.-C., Chen, Y.-Y., Yu, F.-S., Lu, H.-F., Yu, C.-S., Chou, Y.-C., Yu, C.-C., & Chung, J.-G. (2016). Allyl Isothiocyanate Induces Cell Toxicity by Multiple Pathways in Human Breast Cancer Cells. *Http://Dx.Doi.Org/10.1142/S0192415X16500245*, 44(2), 415–437. https://doi.org/10.1142/S0192415X16500245

Bolger, A. M., Lohse, M., & Usadel, B. (2014). Trimmomatic: a flexible trimmer for Illumina sequence data. Bioinformatics, 30(15), 2114–2120. https://doi.org/10.1093/BIOINFORMATICS/BTU170

Borek, V., Morra, M. J., Brown, P. D., & McCaffrey, J. P. (1995). Transformation of the Glucosinolate-Derived Allelochemicals Allyl Isothiocyanate and Allylnitrile in Soil. Journal of Agricultural and Food Chemistry, 43(7), 1935–1940. https://doi.org/10.1021/jf00055a033

Boutoille, D., Corvec, S., Caroff, N., Giraudeau, C., Espaze, E., Caillon, J., Plésiat, P., & Reynaud, A. (2004). Detection of an IS21 insertion sequence in the mexR gene of Pseudomonas aeruginosa increasing β-lactam resistance. FEMS Microbiology Letters, 230(1), 143–146. https://doi.org/10.1016/S0378-1097(03)00882-6

Calmes, B., N’Guyen, G., Dumur, J., Brisach, C. A., Campion, C., Lacomi, B., PignÉ, S., Dias, E., Macherel, D., Guillemette, T., & Simoneau, P. (2015). Glucosinolate-derived isothiocyanates impact mitochondrial function in fungal cells and elicit an oxidative stress response necessary for growth recovery. Frontiers in Plant Science, 6(June), 414. https://doi.org/10.3389/fpls.2015.00414

Chellemi, D. O., Olson, S. M., Mitchell, D. J., Secker, I., & McSorley, R. (1997). Adaptation of soil solarization to the integrated management of soilborne pests of tomato under humid conditions. Phytopathology, 87(3), 250–258. https://doi.org/10.1094/PHYTO.1997.87.3.250

Ciampi-Panno, L., Fernandez, C., Bustamante, P., Andrade, N., Ojeda, S., & Contreras, A. (1989). Biological control of bacterial wilt of potatoes caused byPseudomonas solanacearum. American Potato Journal, 66(5), 315–332. https://doi.org/10.1007/BF02854019

Dufour, V., Stahl, M., Rosenfeld, E., Stintzi, A., & Baysse, C. (2013). Insights into the mode of action of benzyl isothiocyanate on Campylobacter jejuni. Applied and Environmental Microbiology, 79(22), 6958–6968. https://doi.org/10.1128/AEM.01967-13

Elphinstone, J. G. (2005). The current bacterial wilt situation: a global overview. Bacterial Wilt Disease and the Ralstonia Solanacearum Species Complex, 9–28.

Frick, A., Zebarth, B. J., & Szeto, S. Y. (1998). Behavior of the Soil Fumigant Methyl Isothiocyanate in Repacked Soil Columns. Journal of Environmental Quality, 27(5), 1158–1169. https://doi.org/10.2134/jeq1998.00472425002700050022x

Gimsing, A L, & Kirkegaard, J. A. (2006). Glucosinolate and isothiocyanate concentration in soil following incorporation of Brassica biofumigants. Soil Biology & Biochemistry, 38, 2255–2264. https://doi.org/10.1016/j.soilbio.2006.01.024

Gimsing, Anne Louise, & Kirkegaard, J. A. (2009). Glucosinolates and biofumigation: fate of glucosinolates and their hydrolysis products in soil. Phytochemistry Reviews, 8(1), 299– 310. https://doi.org/10.1007/s11101-008-9105-5

Gimsing, Anne Louise, Poulsen, J. L., Pedersen, H. L., & Hansen, H. C. B. (2007). Formation and degradation kinetics of the biofumigant benzyl isothiocyanate in soil. Environmental Science and Technology, 41(12), 4271–4276. https://doi.org/10.1021/es061987t

Gonçalves, O. S., Campos, K. F., Assis, J. C. S. de, Fernandes, A. S., Souza, T. S., Rodrigues, L. G. do C., Queiroz, M. V. de, & Santana, M. F. (2020). Transposable elements contribute to the genome plasticity of Ralstonia solanacearum species complex. Microbial Genomics, 6(5), 1–12. https://doi.org/10.1099/MGEN.0.000374

Greene, N. P., Crow, A., Hughes, C., & Koronakis, V. (2015). Structure of a bacterial toxin-activating acyltransferase. Proceedings of the National Academy of Sciences, 112(23), E3058–E3066. https://doi.org/10.1073/PNAS.1503832112

Guarischi-Sousa, R., Puigvert, M., Coll, N. S., Siri, M. I., Pianzzola, M. J., Valls, M., & Setubal, J. C. (2016). Complete genome sequence of the potato pathogen Ralstonia solanacearum UY031. Standards in Genomic Sciences, 11(1), 7. https://doi.org/10.1186/s40793-016-0131-4

Hanschen, F. S., Bru□, N., N., Brodehl, A., Mewis, I., Schreiner, M., Rohn, S., & Kroh, L. W. (2012). *Characterization of Products from the Reaction of Glucosinolate-Derived Isothiocyanates with Cysteine and Lysine Derivatives Formed in Either Model Systems or Broccoli Sprouts*. https://doi.org/10.1021/jf301718g

Hanschen, F. S., Brüggemann, N., Brodehl, A., Mewis, I., Schreiner, M., Rohn, S., & Kroh, L. W. (2012). Characterization of products from the reaction of glucosinolate-derived isothiocyanates with cysteine and lysine derivatives formed in either model systems or broccoli sprouts. Journal of Agricultural and Food Chemistry, 60(31), 7735–7745. https://doi.org/10.1021/jf301718g

Hanschen, F. S., Yim, B., Winkelmann, T., Smalla, K., & Schreiner, M. (2015). Degradation of Biofumigant Isothiocyanates and Allyl Glucosinolate in Soil and Their Effects on the Microbial Community Composition. PloS One, 10(7), e0132931. https://doi.org/10.1371/journal.pone.0132931

Hartz, T. K., Johnstone, P. R., Miyao, E. M., & Davis, R. M. (2005). Mustard cover crops are ineffective in suppressing soilborne disease or improving processing tomato yield. HortScience, 40(7), 2016–2019. https://doi.org/10.21273/hortsci.40.7.2016

Hawkey, J., Hamidian, M., Wick, R. R., Edwards, D. J., Billman-Jacobe, H., Hall, R. M., & Holt, K. E. (2015). ISMapper: identifying transposase insertion sites in bacterial genomes from short read sequence data. BMC Genomics 2015 16:1, 16(1), 1–11. https://doi.org/10.1186/S12864-015-1860-2

Hawkey, J., Monk, J. M., Billman-Jacobe, H., Palsson, B., & Holt, K. E. (2020). Impact of insertion sequences on convergent evolution of Shigella species. PLOS Genetics, 16(7), e1008931. https://doi.org/10.1371/JOURNAL.PGEN.1008931

Hibbing, M. E., Fuqua, C., Parsek, M. R., & Peterson, S. B. (2010). Bacterial competition: Surviving and thriving in the microbial jungle. In Nature Reviews Microbiology (Vol. 8, Issue 1, pp. 15–25). Nature Publishing Group. https://doi.org/10.1038/nrmicro2259

Hommais, F., Krin, E., Laurent-Winter, C., Soutourina, O., Malpertuy, A., Caer, J.-P. Le, Danchin, A., & Bertin, P. (2001). Large-scale monitoring of pleiotropic regulation of gene expression by the prokaryotic nucleoid-associated protein, H-NS. Molecular Microbiology, 40(1), 20–36. https://doi.org/10.1046/J.1365-2958.2001.02358.X

Hu, P., Hollister, E. B., Somenahally, A. C., Hons, F. M., & Gentry, T. J. (2015). Soil bacterial and fungal communities respond differently to various isothiocyanates added for biofumigation. Frontiers in Microbiology, 5, 729. https://doi.org/10.3389/fmicb.2014.00729

Jeong, E. L., & Timmis, J. N. (2000). Novel insertion sequence elements associated with genetic heterogeneity and phenotype conversion in Ralstonia solanacearum. Journal of Bacteriology, 182(16), 4673–4676. https://doi.org/10.1128/JB.182.16.4673-4676.2000

Ji, P., Momol, M. T., Rich, J. R., Olson, S. M., & Jones, J. B. (2007). Development of an Integrated Approach for Managing Bacterial Wilt and Root-Knot on Tomato Under Field Conditions. Plant Disease, 91(10), 1321–1326. https://doi.org/10.1094/PDIS-91-10-1321

Johnson, D. C., Dean, D. R., Smith, A. D., & Johnson, M. K. (2005). STRUCTURE, FUNCTION, AND FORMATION OF BIOLOGICAL IRON-SULFUR CLUSTERS. *Http://Dx.Doi.Org/10.1146/Annurev.Biochem.74.082803.133518*, 74, 247–281. https://doi.org/10.1146/ANNUREV.BIOCHEM.74.082803.133518

Khokhani, D., Lowe-Power, T. M., Tran, T. M., & Allen, C. (2017). A single regulator mediates strategic switching between attachment/spread and growth/virulence in the plant pathogen Ralstonia solanacearum. MBio, 8(5). https://doi.org/10.1128/MBIO.00895-17

Kirkegaard, J. A., & Matthiessen, J. N. (2005). Developing and refining the biofumigation concept.

Kirkegaard, J. A., & Sarwar, M. (1998). Biofumigation potential of brassicas: I. Variation in glucosinolate profiles of diverse field-grown brassicas. Plant and Soil, 201(1), 71–89. https://doi.org/10.1023/A:1004364713152

Kirkegaard, J. A., Sarwar, M., Wong, P. T. W., Mead, A., Howe, G., & Newell, M. (2000). Field studies on the biofumigation of take-all by Brassica break crops. Australian Journal of Agricultural Research, 51(4), 445–456. https://doi.org/10.1071/AR99106

Kirkegaard, J. A., Wong, P. T. W., & Desmarchelier, J. M. (1996). In vitro suppression of fungal root pathogens of cereals by Brassica tissues. Plant Pathology, 45(3), 593–603. https://doi.org/10.1046/j.1365-3059.1996.d01-143.x

Knöppel, A., Näsvall, J., & Andersson, D. I. (2017). Evolution of antibiotic resistance without antibiotic exposure. Antimicrobial Agents and Chemotherapy, 61(11), 1495–1512. https://doi.org/10.1128/AAC.01495-17

Kozakai, R., Ono, T., Hoshino, S., Takahashi, H., Katsuyama, Y., Sugai, Y., Ozaki, T., Teramoto, K., Teramoto, K., Tanaka, K., Abe, I., Asamizu, S., & Onaka, H. (2020). Acyltransferase that catalyses the condensation of polyketide and peptide moieties of goadvionin hybrid lipopeptides. Nature Chemistry 2020 12:9, 12(9), 869–877. https://doi.org/10.1038/s41557-020-0508-2

Kurt, Ş., Güneş, U., & Soylu, E. M. (2011). *In vitro* and *in vivo* antifungal activity of synthetic pure isothiocyanates against *Sclerotinia sclerotiorum*. Pest Management Science, 67(7), 869–875. https://doi.org/10.1002/ps.2126

Lacour, S., Bechet, E., Cozzone, A. J., Mijakovic, I., & Grangeasse, C. (2008). Tyrosine Phosphorylation of the UDP-Glucose Dehydrogenase of Escherichia coli Is at the Crossroads of Colanic Acid Synthesis and Polymyxin Resistance. PLoS ONE, 3(8), e3053. https://doi.org/10.1371/journal.pone.0003053

Larkin, R. P., & Griffin, T. S. (2007). Control of soilborne potato diseases using Brassica green manures. Crop Protection, 26(7), 1067–1077. https://doi.org/10.1016/j.cropro.2006.10.004

Larkin, R. P., & Halloran, J. M. (2015). Management effects of disease-suppressive rotation crops on potato yield and soilborne disease and their economic implications in potato production. American Journal of Potato Research, 91(5), 429–439. https://doi.org/10.1007/s12230-014-9366-z

Li, E., de Jonge, R., Liu, C., Jiang, H., Friman, V. P., Pieterse, C. M. J., Bakker, P. A. H. M., & Jousset, A. (2020). Rapid evolution of bacterial mutualism in the plant rhizosphere. In bioRxiv (p. 2020.12.07.414607). bioRxiv. https://doi.org/10.1101/2020.12.07.414607

Li, Z., Tang, Y., Wu, Y., Zhao, S., Bao, J., Luo, Y., & Li, D. (2017). Structural insights into the committed step of bacterial phospholipid biosynthesis. Nature Communications 2017 8:1, 8(1), 1–14. https://doi.org/10.1038/s41467-017-01821-9

Lin, C.-M., Preston Iii, J. F., & Wei, C.-I. (2000). Antibacterial Mechanism of Allyl Isothiocyanate. Journal of Food Protection, 63(6), 727–734. http://jfoodprotection.org/doi/pdf/10.4315/0362-028X-63.6.727

Loewe, L., Textor, V., & Scherer, S. (2003). High Deleterious Genomic Mutation Rate in Stationary Phase of Escherichia coli. Science, 302(5650), 1558–1560. https://doi.org/10.1126/science.1087911

Lord, J. S., Lazzeri, L., Atkinson, H. J., & Urwin, P. E. (2011). Biofumigation for Control of Pale Potato Cyst Nematodes: Activity of Brassica Leaf Extracts and Green Manures on Globodera pallida in Vitro and in Soil. J. Agric. Food Chem, 59, 7882–7890. https://doi.org/10.1021/jf200925k

Luciano, F. B., & Holley, R. A. (2009). Enzymatic inhibition by allyl isothiocyanate and factors affecting its antimicrobial action against Escherichia coli O157:H7. International Journal of Food Microbiology, 131(2–3), 240–245. https://doi.org/10.1016/j.ijfoodmicro.2009.03.005

Manici, L. M., Lazzeri, L., & Palmieri, S. (1997). In Vitro Fungitoxic Activity of Some Glucosinolates and Their Enzyme-Derived Products toward Plant Pathogenic Fungi. Journal of Agricultural and Food Chemistry, 45(7), 2768–2773. https://doi.org/10.1021/jf9608635

Marshall, C. G., Zolli, M., & Wright, G. D. (1999). Molecular mechanism of VanHst, an α ketoacid dehydrogenase required for glycopeptide antibiotic resistance from a glycopeptide producing organism. Biochemistry, 38(26), 8485–8491. https://doi.org/10.1021/bi982843x

Matthiessen, J. N., & Kirkegaard, J. A. (2006). Biofumigation and Enhanced Biodegradation: Opportunity and Challenge in Soilborne Pest and Disease Management. Critical Reviews in Plant Sciences, 25(3), 235–265. https://doi.org/10.1080/07352680600611543

Matthiessen, J. N., & Shackleton, M. A. (2005). Biofumigation: environmental impacts on the biological activity of diverse pure and plant-derived isothiocyanates. Pest Management Science, 61(11), 1043–1051. https://doi.org/10.1002/ps.1086

Mazzola, M., & Gu, Y. H. (2002). Wheat genotype-specific induction of soil microbial communities suppressive to disease incited by Rhizoctonia solani Anastomosis Group (AG)-5 and AG-8. Phytopathology, 92(12), 1300–1307. https://doi.org/10.1094/PHYTO.2002.92.12.1300

Mazzola, M., Hewavitharana, S. S., & Strauss, S. L. (2015). *Brassica* Seed Meal Soil Amendments Transform the Rhizosphere Microbiome and Improve Apple Production Through Resistance to Pathogen Reinfestation. Phytopathology, 105(4), 460–469. https://doi.org/10.1094/PHYTO-09-14-0247-R

Mondal, M. F., Asaduzzaman, M., & Asao, T. (2015). Adverse Effects of Allelopathy from Legume Crops and Its Possible Avoidance. American Journal of Plant Sciences, 06(06), 804–810. https://doi.org/10.4236/ajps.2015.66086

Murakami, K., Ono, T., Viducic, D., Kayama, S., Mori, M., Hirota, K., Nemoto, K., & Miyake, Y. (2005). Role for rpoS gene of Pseudomonas aeruginosa in antibiotic tolerance. FEMS Microbiology Letters, 242(1), 161–167. https://doi.org/10.1016/j.femsle.2004.11.005

Navarro Llorens, J. M., Tormo, A., & Martínez-García, E. (2010). Stationary phase in gram-negative bacteria. In FEMS Microbiology Reviews (Vol. 34, Issue 4, pp. 476–495). Blackwell Publishing Ltd. https://doi.org/10.1111/j.1574-6976.2010.00213.x

Neubauer, C., Heitmann, B., & Müller, C. (2014). Biofumigation potential of Brassicaceae cultivars to Verticillium dahliae. European Journal of Plant Pathology, 140(2), 341–352. https://doi.org/10.1007/s10658-014-0467-9

Ngala, B. M., Woods, S. R., & Back, M. A. (2015). Sinigrin degradation and G. pallida suppression in soil cultivated with brassicas under controlled environmental conditions. Applied Soil Ecology, 95, 9–14. https://doi.org/10.1016/j.apsoil.2015.05.009

Ohlrogge, J., & Browse, J. (1995). Lipid biosynthesis. The Plant Cell, 7(7), 957. https://doi.org/10.1105/TPC.7.7.957

Olivier, C., Vaughn, S. F., Mizubuti, E. S. G., & Loria, R. (1999). Variation in allyl isothiocyanate production within Brassica species and correlation with fungicidal activity. Journal of Chemical Ecology, 25(12), 2687–2701. https://doi.org/10.1023/A:1020895306588

Olsen, I. (2015). Biofilm-specific antibiotic tolerance and resistance. In European Journal of Clinical Microbiology and Infectious Diseases (Vol. 34, Issue 5, pp. 877–886). Springer Verlag. https://doi.org/10.1007/s10096-015-2323-z

Palmer, T., Sargent, F., & Berks, B. C. (2005). Export of complex cofactor-containing proteins by the bacterial Tat pathway. Trends in Microbiology, 13(4), 175–180. https://doi.org/10.1016/J.TIM.2005.02.002

Pasternak, C., Dulermo, R., Ton-Hoang, B., Debuchy, R., Siguier, P., Coste, G., Chandler, M., & Sommer, S. (2013). IS *Dra 2* transposition in *D einococcus radiodurans* is downregulated by TnpB. Molecular Microbiology, 88(2), 443–455. https://doi.org/10.1111/mmi.12194

Perrier, A., Barlet, X., Rengel, D., Prior, P., Poussier, S., Genin, S., & Guidot, A. (2019). Spontaneous mutations in a regulatory gene induce phenotypic heterogeneity and adaptation of Ralstonia solanacearum to changing environments. Environmental Microbiology, 21(8), 3140–3152. https://doi.org/10.1111/1462-2920.14717

Poirel, L., Decousser, J. W., & Nordmann, P. (2003). Insertion sequence ISEcp1B is involved in expression and mobilization of a blaCTX-M β-lactamase gene. Antimicrobial Agents and Chemotherapy, 47(9), 2938–2945. https://doi.org/10.1128/AAC.47.9.2938-2945.2003

Qin, S., Gan, J., Liu, W., & Becker, J. O. (2004). *Degradation and Adsorption of Fosthiazate in Soil*. https://doi.org/10.1021/jf049094c

Ramesh, R., Joshi, A. A., & Ghanekar, M. P. (2009). Pseudomonads: major antagonistic endophytic bacteria to suppress bacterial wilt pathogen, Ralstonia solanacearum in the eggplant (Solanum melongena L.). World Journal of Microbiology and Biotechnology, 25(1), 47–55. https://doi.org/10.1007/s11274-008-9859-3

Ranjan, V. K., Mukherjee, S., Thakur, S., Gupta, K., & Chakraborty, R. (2021). Pandrug-resistant Pseudomonas sp. expresses New Delhi metallo-β-lactamase-1 and consumes ampicillin as sole carbon source. Clinical Microbiology and Infection, 27(3), 472.e1-472.e5. https://doi.org/10.1016/j.cmi.2020.10.032

Rodionova, I. A., Zhang, Z., Aboulwafa, M., & Saier, M. H. (2020). UDP-glucose dehydrogenase Ugd in E. coli is activated by Gmd and RffD, is inhibited by CheY, and regulates swarming. In bioRxiv (p. 2020.01.08.899336). bioRxiv. https://doi.org/10.1101/2020.01.08.899336

Rudolph, R. E., Sams, C., Steiner, R., Thomas, S. H., Walker, S., & Uchanski, M. E. (2015). Biofumigation performance of four brassica crops in a green chile pepper (Capsicum annuum) rotation system in southern New Mexico. HortScience, 50(2), 247–253. https://doi.org/10.21273/hortsci.50.2.247

Rumberger, A., & Marschner, P. (2003). 2-Phenylethylisothiocyanate concentration and microbial community composition in the rhizosphere of canola. Soil Biology and Biochemistry, 35(3), 445–452. https://doi.org/10.1016/S0038-0717(02)00296-1

Salanoubat, M., Genin, S., Artiguenave, F., Gouzy, J., Mangenot, S., Arlat, M., Billault, A., Brottier, P., Camus, J. C., Cattolico, L., Chandler, M., Choisne, N., Claudel-Renard, C., Cunnac, S., Demange, N., Gaspin, C., Lavie, M., Moisan, A., Robert, C., … Boucher, C. A. (2002). Genome sequence of the plant pathogen Ralstonia solanacearum. Nature, 415(6871), 497–502. https://doi.org/10.1038/415497a

Sarwar, M., Kirkegaard, J. A., Wong, P. T. W., & Desmarchelier, J. M. (1998). Biofumigation potential of brassicas. Plant and Soil, 201(1), 103–112. https://doi.org/10.1023/A:1004381129991

Savary, S., Ficke, A., Aubertot, J. N., & Hollier, C. (2012). Crop losses due to diseases and their implications for global food production losses and food security. In Food Security (Vol. 4, Issue 4, pp. 519–537). Springer. https://doi.org/10.1007/s12571-012-0200-5

Seemann, T. (2014). Prokka: rapid prokaryotic genome annotation. Bioinformatics, 30(14), 2068–2069. https://doi.org/10.1093/bioinformatics/btu153

Seemann, T. (2015). *Snippy: fast bacterial variant calling from NGS reads.* https://github.com/tseemann/snippy.

Shigemizu, D., Miya, F., Akiyama, S., Okuda, S., Boroevich, K. A., Fujimoto, A., Nakagawa, H., Ozaki, K., Niida, S., Kanemura, Y., Okamoto, N., Saitoh, S., Kato, M., Yamasaki, M., Matsunaga, T., Mutai, H., Kosaki, K., & Tsunoda, T. (2018). IMSindel: An accurate intermediate-size indel detection tool incorporating de novo assembly and gapped global-local alignment with split read analysis. Scientific Reports 2018 8:1, 8(1), 1–9. https://doi.org/10.1038/s41598-018-23978-z

Smith, B. J., & Kirkegaard, J. A. (2002). In vitro inhibition of soil microorganisms by 2-phenylethyl isothiocyanate. Plant Pathology, 51(5), 585–593. https://doi.org/10.1046/j.1365-3059.2002.00744.x

Sofrata, A., Santangelo, E. M., Azeem, M., Borg-Karlson, A.-K., Gustafsson, A., & Pütsep, K. (2011). Benzyl Isothiocyanate, a Major Component from the Roots of Salvadora Persica Is Highly Active against Gram-Negative Bacteria. PLoS ONE, 6(8), e23045. https://doi.org/10.1371/journal.pone.0023045

Stirling, G. R., & Stirling, A. M. (2003). The potential of Brassica green manure crops for controlling root-knot nematode (Meloidogyne javanica) on horticultural crops in a subtropical environment. Australian Journal of Experimental Agriculture, 43(6), 623– 630. https://doi.org/10.1071/EA02175

Stringlis, I. A., Yu, K., Feussner, K., De Jonge, R., Van Bentum, S., Van Verk, M. C., Berendsen, R. L., Bakker, P. A. H. M., Feussner, I., & Pieterse, C. M. J. (2018). MYB72-dependent coumarin exudation shapes root microbiome assembly to promote plant health. Proceedings of the National Academy of Sciences of the United States of America, 115(22), E5213–E5222. https://doi.org/10.1073/pnas.1722335115

Tatusova, T., DiCuccio, M., Badretdin, A., Chetvernin, V., Nawrocki, E. P., Zaslavsky, L., Lomsadze, A., Pruitt, K. D., Borodovsky, M., & Ostell, J. (2016). NCBI prokaryotic genome annotation pipeline. Nucleic Acids Research, 44(14), 6614–6624. https://doi.org/10.1093/NAR/GKW569

Tusskorn, O., Senggunprai, L., Prawan, A., Kukongviriyapan, U., & Kukongviriyapan, V. (2013). Phenethyl isothiocyanate induces calcium mobilization and mitochondrial cell death pathway in cholangiocarcinoma KKU-M214 cells. BMC Cancer 2013 13:1, 13(1), 1–12. https://doi.org/10.1186/1471-2407-13-571

Vaughn, S. F., Palmquist, D. E., Duval, S. M., & Berhow, M. A. (n.d.). Herbicidal Activity of Glucosinolate-Containing Seedmeals (Vol. 54, Issue 4).

Vervoort, M. T. W., Vonk, J. A., Brolsma, K. M., Schütze, W., Quist, C. W., De Goede, R. G. M., Hoffland, E., Bakker, J., Mulder, C., Hallmann, J., & Helder, J. (2014). Release of isothiocyanates does not explain the effects of biofumigation with Indian mustard cultivars on nematode assemblages. Soil Biology and Biochemistry, 68, 200–207. https://doi.org/10.1016/j.soilbio.2013.10.008

Wang, D., Rosen, C., Kinkel, L., Cao, A., Tharayil, N., & Gerik, J. (2009). Production of methyl sulfide and dimethyl disulfide from soil-incorporated plant materials and implications for controlling soilborne pathogens. Plant and Soil, 324(1–2), 185–197. https://doi.org/10.1007/s11104-009-9943-y

Warton, B., Matthiessen, J. N., & Shackleton, M. A. (2001). Glucosinolate content and isothiocyanate evolution - Two measures of the biofumigation potential of plants. Journal of Agricultural and Food Chemistry, 49(11), 5244–5250. https://doi.org/10.1021/jf010545s

Wick, R. R., Judd, L. M., Gorrie, C. L., & Holt, K. E. (2017). Unicycler: Resolving bacterial genome assemblies from short and long sequencing reads. PLOS Computational Biology, 13(6), e1005595. https://doi.org/10.1371/JOURNAL.PCBI.1005595

Wink, M. (2013). Evolution of secondary metabolites in legumes (Fabaceae). South African Journal of Botany, 89, 164–175. https://doi.org/10.1016/j.sajb.2013.06.006

Xie, Z., & Tang, H. (2017). ISEScan: automated identification of insertion sequence elements in prokaryotic genomes. Bioinformatics, 33(21), 3340–3347. https://doi.org/10.1093/BIOINFORMATICS/BTX433

Yabuuchi, E., Kosako, Y., Yano, I., Hotta, H., & Nishiuchi, Y. (1995). Transfer of Two *Burkholderia* and An *Alcaligenes* Species to *Ralstonia* Gen. Nov. Microbiology and Immunology, 39(11), 897–904. https://doi.org/10.1111/j.1348-0421.1995.tb03275.x

Yim, B., Hanschen, F. S., Wrede, A., Nitt, H., Schreiner, M., Smalla, K., & Winkelmann, T. (2016). Effects of biofumigation using Brassica juncea and Raphanus sativus in comparison to disinfection using Basamid on apple plant growth and soil microbial communities at three field sites with replant disease. Plant and Soil, 406(1–2), 389– 408. https://doi.org/10.1007/s11104-016-2876-3

Yun, Y.-K., Kim, H.-K., Kim, J.-R., Hwang, K., & Ahn, Y.-J. (2012). Contact and fumigant toxicity of *Armoracia rusticana* essential oil, allyl isothiocyanate and related compounds to *Dermatophagoides farinae*. Pest Management Science, 68(5), 788–794. https://doi.org/10.1002/ps.2327

Zampieri, M., Enke, T., Chubukov, V., Ricci, V., Piddock, L., & Sauer, U. (2017). Metabolic constraints on the evolution of antibiotic resistance. Molecular Systems Biology, 13(3), 917. https://doi.org/10.15252/msb.20167028

